# *Helicobacter pylori* genome aggregation database reveals complex evolutionary forces shaping its genomic landscape and clinical impact in East Asia

**DOI:** 10.1101/2025.08.13.669870

**Authors:** Lintao Luo, Xiaoqiong Tang, Hao Chen, Yunhui Liu, Limei Zhang, Jie Zhong, Bowen Li, Yutong Jiang, Zihao Zhu, Yuzhu Wang, Yuguo Huang, Fengxiao Bu, Huijun Yuan, Chao Liu, Haohao Dong, Binwu Ying, Hong Li, Renkuan Tang, Mengge Wang, Guanglin He

## Abstract

We assembled HPgnomAD, a comprehensive global genome aggregation database of *Helicobacter pylori* (*H. pylori*) featuring 7,544 high-quality genomes from 278 populations, along with the first species-wide haplotype reference panel for genotype imputation. The panel provides high accuracy across diverse *H. pylori* populations, including those from low-coverage data, and is openly accessible with integrated analysis tools (https://www.hpgnomad.top/). Variant discovery revealed 1.82 million SNPs and 0.65 million InDels, with African strains exhibiting the most tremendous diversity and East Asian strains showing unexpectedly high novel variation. Fine-scale analysis of the hpEAsia lineage revealed six new sublineages shaped by altitude-, latitude-related divergence and region-specific admixture. Phylogenomic dating revealed two divergence waves in East Asia, paralleling Upper Paleolithic settlement and Neolithic human expansions, with highland–lowland separation at ∼18.8 kya, followed by the formation of complex geography-related population substructures. Genome-wide scans revealed adaptive loci related to metal acquisition, nitrogen metabolism, surface adhesion, and membrane transport, including altitude-associated highly differentiated variants linked to antibiotic resistance and gastric disease. By integrating genomic, evolutionary, environmental, and clinical data, HPgnomAD offers a framework for understanding *H. pylori* evolution, host‒pathogen adaptation, and precision medicine worldwide.

## Introduction

*Helicobacter pylori* (*H. pylori*) predominantly colonizes the human stomach and has coexisted with humans for approximately 70,000 years; *H. pylori* is now known as a primary etiological agent of chronic inflammation, gastritis, peptic ulcers, and gastric malignancies ^1–8^. Its genome consists of a circular, double-stranded DNA molecule approximately 1.6–1.7 million base pairs in length ^9^. Extensive recombination and high mutation rates have driven exceptional genetic diversity, nearly 50 times greater than that observed in humans ^10,11^. As a result, *H. pylori* strains display considerable genotypic and phenotypic variation across global populations ^12–16^. Because *H. pylori* evolution closely mirrors human evolutionary history, its genome serves as a powerful proxy for reconstructing ancient human migrations ^14,17,18^. Large-scale human genomic data and ancient DNA studies further suggest a more complex evolutionary history for global *H. pylori* populations than previously assumed ^19–23^. Recent genomic studies have significantly advanced our understanding of the global distribution and phylogeographic structure of *H. pylori*, particularly through efforts such as the *H. pylori* Genome Project (HpGP), which systematically classified strains into distinct populations and subpopulations ^15^. Building on this paradigm and a larger-scale *H. pylori* genomic dataset, Tourrette *et al*. introduced the concept of “ecospecies” to reconcile discrepancies between genomic divergence and species definitions, incorporating large-scale data, including Indigenous American strains ^16^. While such studies have generated invaluable resources for exploring *H. pylori* evolutionary dynamics, high-quality, population-wide databases capturing SNPs, InDels, and structural variants remain scarce.

Previous genetic studies have characterized the global population structure of *H. pylori*, revealing distinct clustering patterns and subpopulations ^15,24^. In East Asia, lineages such as hpEAsia, hpNorthAsia, and hpAsia2 appear to contribute significantly to regional population dynamics ^15,24^. Clinical epidemiological data have further highlighted the high prevalence of gastric cancer in East Asia, and with a differentiated occurrence rate among both highland and lowland East Asian populations ^25–28^. Recently, the expansion of genomic resources from East Asian *H. pylori* strains has enabled a deeper investigation into their diversity and potential roles in the pathogenesis of gastrointestinal disease ^29,30^. You *et al*. identified seven subregional clusters by analyzing 381 genomes and reported pronounced allelic variation at several loci, potentially linked to ecological adaptation ^29^. In parallel, Suzuki *et al*. analyzed 88 genomes from Okinawa, including 21 newly sequenced strains, to support a model of Paleolithic human migration from Africa to East Asia ^30^. Despite these advances, both studies present notable limitations in terms of the availability of high-quality databases, the scale of population stratification, and deep demographic reconstruction from a worldwide perspective. Most works are restricted to *H. pylori* isolate datasets, limiting the ability to draw continent-wide demographic and temporal inferences. Suzuki *et al*. relied on multi-locus sequence typing (MLST), which excludes genome-wide variation, potentially missing subtle yet biologically essential signals. While large-scale resources from the studies of HpGP and Elise *et al.* include East Asian *H. pylori* genomes ^15,16^, they provide limited insight into high-resolution genetic variation, fine-scale structure, biological adaptation, and historical relevance. Moreover, population-specific reference datasets for clinical and basic research remain insufficiently developed. The prevalence of *H. pylori* infection varies markedly across East Asia and is shaped by complex evolutionary pressures. These include extreme conditions, such as cold, hypoxia, and resource scarcity on the Qinghai-Xizang Plateau; distinct dietary practices across pastoral-agricultural regions of Northeast Asia; seafood-rich diets in southern East Asia; and heterogeneous pathogen exposures ^31–35^. Large-scale human genomic studies have identified adaptive signatures driven by these pressures ^32,36,37^. Additionally, human genetic modeling based on autosomal, Y-chromosomal, and mitochondrial data has revealed detailed population substructures linked to geography or language in East Asia and other regions ^19,21,33,38^. However, how complex human migration and admixture events affect the diversity and structure of geographically distinct *H. pylori* populations and the extent to which such evolutionary factors shape *H. pylori*’s genomic diversity and contribute to region-specific clinical outcomes remain unclear.

To address the limitations posed by the absence of a comprehensive genomic database for *H. pylori* and the scarcity of prior genomic diversity initiatives, we assembled a dataset of 8,010 high-quality *H. pylori* whole-genome sequences from both in-house sources and published studies ^12,15,29,39–62^. Using this dataset, we constructed the *H. pylori* Genome Aggregation Database (HPgnomAD) to investigate genomic diversity and variant spectra across diverse *H. pylori* lineages ^12,15,29,39–62^ (**Fig. 1a; Extended Data Fig. 1, and Extended Data Table 1**). We employed multiple state-of-the-art population admixture modeling approaches, including model-based ADMIXTURE, fineSTRUCTURE, ChromoPainter, and BEAST-based phylogenetic reconstruction, to resolve the fine-scale population structure and infer the divergence times and expansion patterns of genetically distinct *H. pylori* lineages. In parallel, we applied various statistical methods (genome-wide association studies (GWAS), *F*_ST_, and *dN/dS* analyses) to detect signals of natural selection among highland and lowland sublineages, as well as between northern and southern East Asian populations. These analyses revealed adaptive signatures with clinically relevant implications for phenotypic variation. To facilitate broader access, we developed an online platform (https://www.hpgnomad.top/) that catalogs the observed genomic variants, polymorphisms, and associated clinical features of *H. pylori*, providing a resource for researchers and clinicians. By integrating evolutionary and medical perspectives, we present a systematic characterization of the genomic architecture and adaptive trajectories of *H. pylori* in East Asia. Collectively, these findings refine our understanding of host–pathogen co-evolution and offer insights into population-specific prevention and control strategies.

**Fig. 1.**
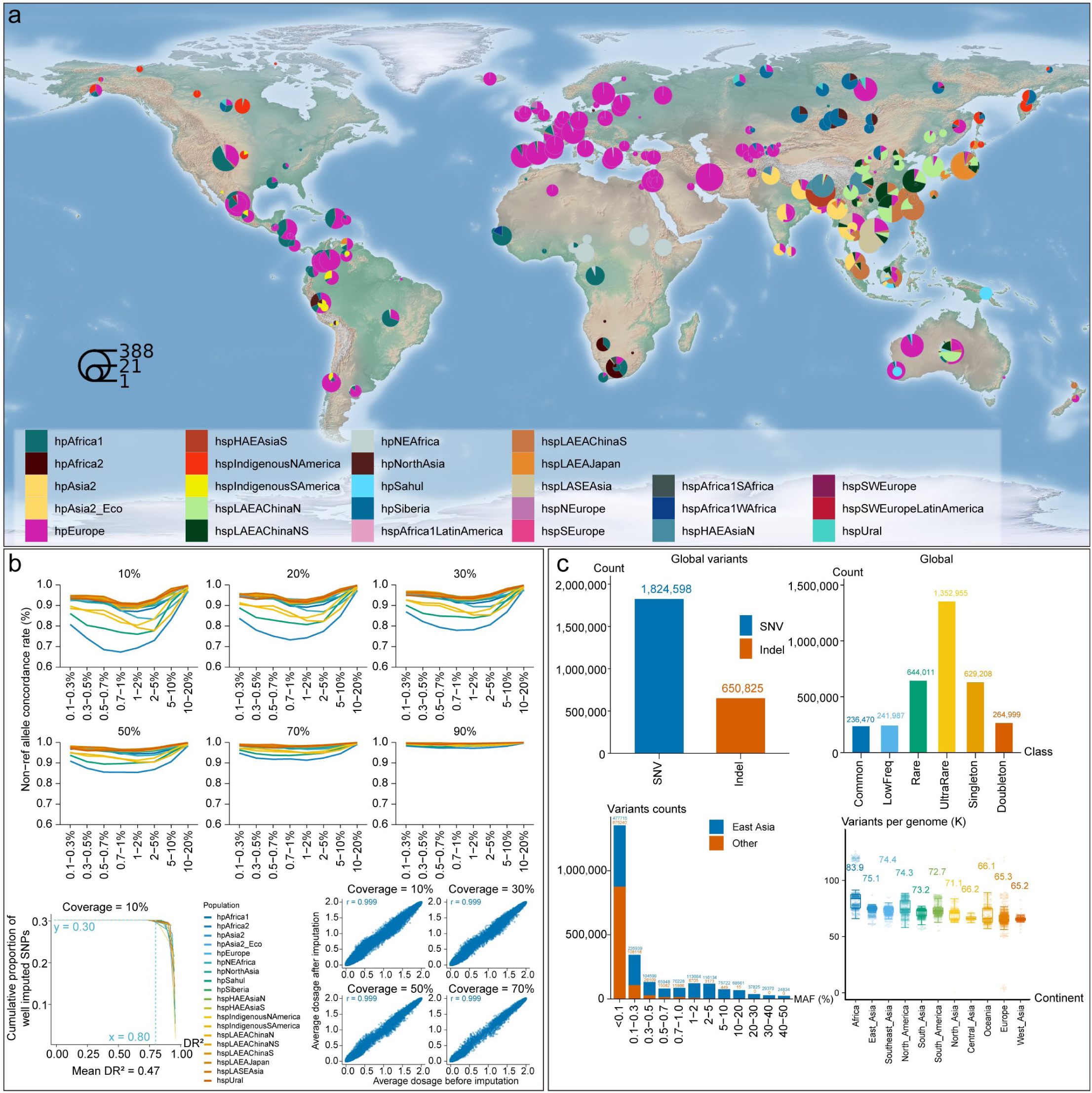
Sample distribution in the HPgnomAD database, performance evaluation of haplotype imputation reference panel , and variant discovery. **a**, Geographical distribution of the samples used in this study. The colors indicate strain classifications, and the marker sizes correspond to the sample sizes. **b**, Reference panel performance evaluation. Upper panel: Genotype concordance by minor allele frequency (MAF) bin for each population, simulated at sequencing coverages of 10%, 20%, 30%, 50%, 70% and 90%. Lower left: cumulative distribution function of dosage R^2^ (DR^2^) at 10% coverage; the x-axis shows the DR^2^ threshold, and the y-axis shows the fraction of total imputed SNPs with DR^2^. Lower right panel: Mean dosage before versus after imputation across coverage levels; Pearson correlation coefficients (r) are indicated. c, Variant discovery in the HPgnomAD database. Variants are classified by common (MAF ≥ 5%); low-frequency (1% ≤ MAF < 5%); rare (0.1% ≤ MAF < 1%); ultra-rare (MAF < 0.1%); doubletons (variants observed in two samples); and singletons (variants observed in a single sample).

## Results

### HPgnomAD construction, haplotype reference panel reconstruction, and variant discovery

We compiled high-quality *H. pylori* whole-genome sequences from in-house resources, published studies ^15,16,29,30,51,58^, and public repositories, including EnteroBase and the National Library of Medicine. This effort produced the HPgnomAD resource, comprising 8,010 genomes from 278 global sampling sites. After stringent quality control, we curated a robust dataset of 7,544 genomes (7,542 *H. pylori* and 2 *H. acinonychis*) for foundational population genetic and evolutionary research (**Fig. 1a; Extended Data Fig. 1a**). From HPgnomAD, we constructed an *H. pylori* haplotype reference panel and assessed its imputation performance (**Fig. 1b; Extended Data Fig. 1b**). Simulations across varying sequencing coverages revealed strong concordance for non-reference alleles, with accuracy increasing at higher coverage. At 10% coverage, agreement exceeded 60% across all populations and minor allele frequency (MAF) bins. Under these conditions, the mean dosage R² (DR²) was 0.47, with 34.1% of the 17,109,424 imputed sites exceeding DR² > 0.8; applying a DR² ≥ 0.8 threshold retained ∼30% of the sites as high quality. The cumulative DR² distributions were consistent across populations, and the Pearson correlations between the mean dosages before and after imputation remained high (r = 0.99) at all coverages. Collectively, these results demonstrate that the haplotype imputation reference panel enables accurate and consistent imputation across diverse *H. pylori* populations, even at low sequencing depths. The panel is publicly available through HPgnomAD (https://www.hpgnomad.top), where users can upload local data or download reference panels for imputation.

Using 7,544 high-quality *H. pylori* genomes, we characterized the distribution of SNPs and InDels. Overall, we found 1,824,598 SNPs and 650,825 InDels (**Fig. 1c**). These variants include 236,470 common, 241,987 low-frequency, 644,011 rare, and 1,352,955 ultra-rare sites, the latter comprising 629,208 singletons and 264,999 doubletons. Geographical analysis revealed that African strains have the greatest number of variants per genome (average: ∼83.9 × 10^3^), reflecting a long evolutionary history and ancient origin. East Asian strains ranked second (average: ∼75.1 × 10^3^), indicating unexpectedly high genetic diversity in this region. Among the 1,657 East Asian genomes, we identified 161,974 common, 222,578 low-frequency, 600,358 rare, and 427,723 ultra-rare variants (**Extended Data Fig. 1c**). The East Asian samples included nearly all the high-frequency sites (MAF ≥ 20%), whereas the MAF < 0.1% category alone contained 1,352,955 sites, which was the most considerable portion of the variants (**Fig. 1c**). Variant counts decrease sharply as the MAF increases, leaving only 24,834 sites in the 40∼50% range. Low-frequency variants are likely recent in origin, influenced by region-specific selective pressures and demographic history, and may be associated with distinct clinical pathogenicity; thus, they merit detailed study. In East Asia, 1,518 of the 1,657 strains (91.61%) belong to the hpEAsia population (**Extended Data Fig. 1c**). However, this broad classification does not capture the complex genetic structure of these strains, highlighting the need for more refined classification.

### Fine-scale genetic substructure and complex genomic diversity of hpEAsia strains

To explore the global genetic landscape of *H. pylori* strains, we first conducted principal component analysis (PCA) and uniform manifold approximation and projection (UMAP) using the core genomes of 7,544 strains (**Fig. 2a and Extended Data Fig. 2**). The overall patterns of *H. pylori* strains worldwide aligned with those recently described via HpGP, but more complex patterns appeared in East Asian strains because of the large sample size. We observed that although most East Asian *H. pylori* strains formed a distinct cluster, a subset showed a continuous gradient along PC1 that overlapped with some European strains (**Fig. 2a**). This pattern indicates historical mixing across Eurasia. When PCA points were colored based on genetic classification rather than geographic origin (**Extended Data Fig. 2**), hpEAsia strains again formed a separate cluster, distinct from other lineages along PC1. Conversely, hpAsia2, which is mainly found in mainland Southeast Asia and South Asia, remained genetically separate along PC2 despite being geographically close to East Asian strains, suggesting an independent evolutionary path.

**Fig. 2.**
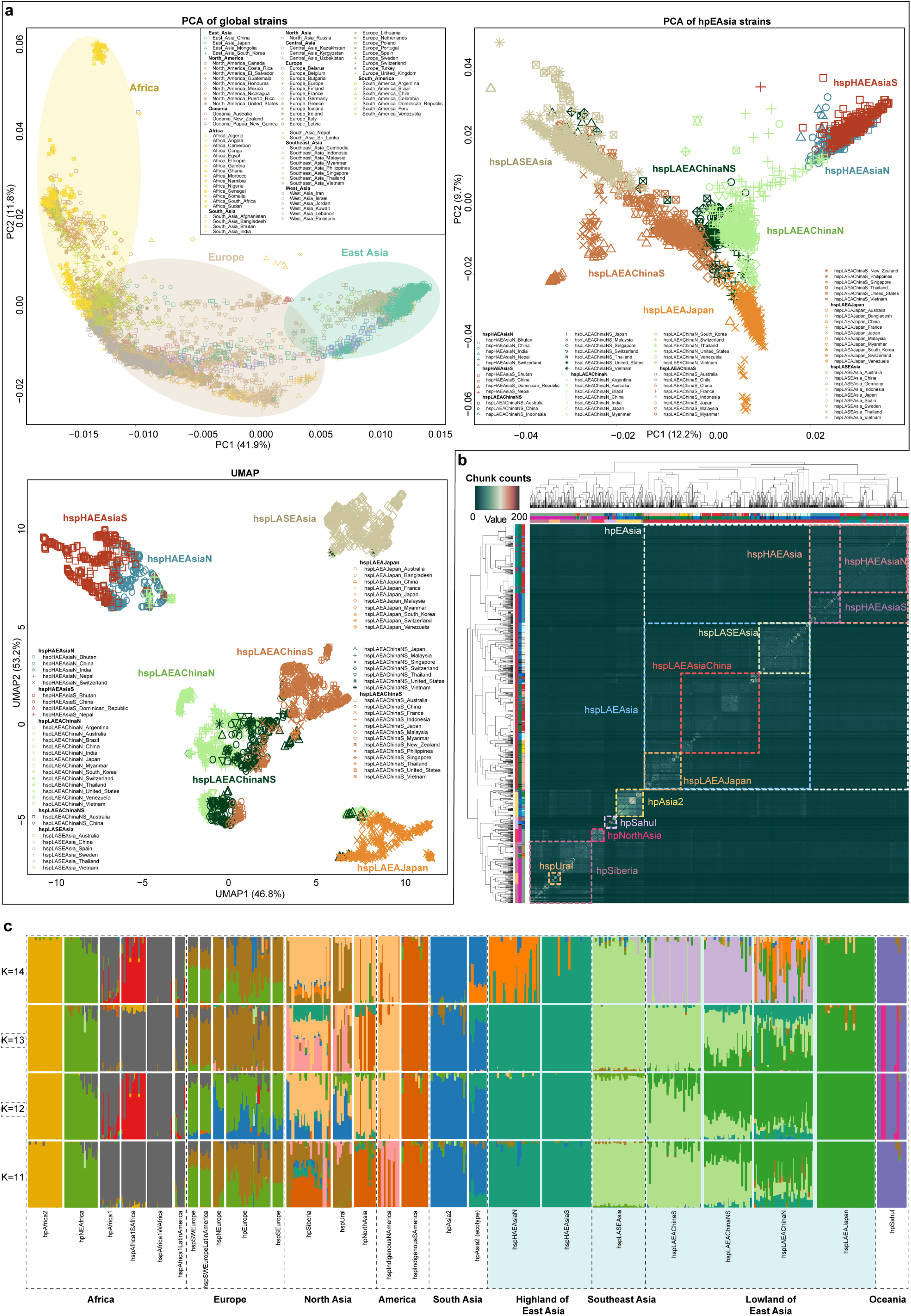
Genetic structure of global *H. pylori* strains. **a**, PCA of global and hpEAsia strains and UMAP of hpEAsia strains. For the global PCA, points are colored according to the sampling continent and shaped by the sampling country. For both the hpEAsia PCA and UMAP, the points are colored according to genetic background (see **Extended Data Fig. 2**). **b,** FineSTRUCTURE analysis of *H. pylori* strains (see **Extended Data Fig. 4** for more details). **c,** ADMIXTURE analysis showing the genetic structure of strains at K = 11–14 (see **Extended Data Table 2** for more details). The optimal K values, as determined by cross-validation, were 12 and 13. Shaded areas highlight hpEAsia strains.

Phylogenetic reconstruction revealed two distinct clades within hpAsia2. Genome-wide association testing, pairwise *F*_ST_ estimates, and analyses of genomic architecture collectively indicated that this clade represents an ecologically distinct population, hereafter called the hpAsia2 ecotype (**Extended Data Fig. 3**). For the hpEAsia strains, we used ordinary least squares regression on the strains from the East Asia subset to examine the relationships between geographic variables and the first three principal components (PCs) (PC1, PC2, and PC3). Each component was standardized through z-score transformation. Altitude was strongly positively correlated with PC1 (R = 0.502, *p* < 0.001), indicating clear differentiation between high- and low-altitude populations. Latitude was the most significant predictor of PC2 (R = 0.178, *p* < 0.001), suggesting that PC2 mainly captures north–south divergence (**Extended Data Table 2**). These results demonstrate that the genomic diversity and structure of East Asian *H. pylori* are closely influenced by geographic factors, especially altitude and latitude, emphasizing the key role of environmental variables, particularly altitude, in shaping population structure.

Furthermore, to characterize the fine-scale population structure of the hpEAsia group in our HPgnomAD dataset, we used fineSTRUCTURE to compute the coancestry matrix (**Fig. 2b and Extended Data Fig. 4**). The analysis revealed a complex demographic landscape among globally spread *H. pylori* lineages and significant diversity within the previously defined hpEAsia subpopulation. Following established naming conventions, we classified clusters by altitude (H/L) and latitude (N/S) within geographic regions. At the primary level, hpEAsia is split into hspHAEAsia (high-altitude) and hspLAEAsia (low-altitude) lineages. hspHAEAsia is subdivided into hspHAEAsiaN (northern Qinghai–Tibet Plateau) and hspHAEAsiaS (southern Himalayas, notably Bhutan), whereas hspLAEAsia is split into hspLASEAsia (Vietnam, Thailand), hspLAEAChina, and hspLAEAJapan. hspLAEAChina further separated into hspLAEAChinaS (southern China), hspLAEAChinaN (northern China), and a hybrid central cluster (hspLAEAChinaNS) enriched in admixed strains from the north–south transition zone.

To better understand the ancestral makeup of the hpEAsia subclusters, we performed a model-based ADMIXTURE analysis with a comprehensive global strain panel to reduce bias. Cross-validation revealed that K = 13 was the best fit for capturing the primary continental ancestry components that shape the admixture patterns. We observed meaningful genetic differences between East Asian *H. pylori* genomes and those from nearby regions such as North Asia, South Asia, and the Americas. In the most suitable model, three major components shaped hpEAsia diversity (**Fig. 2c**): a green component enriched in highland Lhasa and Bhutan, a light green component common in Southeast Asia, and a deep green component dominant in Japan, mirroring fineSTRUCTURE clusters. Chinese lineages presented varying admixture from all three lineages, with hspLAEAChinaNS exhibiting an intermediate profile between hspLAEAChinaS and hspLAEAChinaN. Increasing K to 14 separated hspHAEAsiaN and hspHAEAsiaS into distinct components, indicating unique genetic profiles and admixture histories. The concordance among ADMIXTURE, fineSTRUCTURE, and PCA underscores the presence of fine-scale, geographically structured genetic signatures within hpEAsia.

### Deep evolutionary history of *H. pylori* indicates a temporal link between strain divergence and human migration

Previous genomic studies have reconstructed complex human migration patterns, including out-of-Africa dispersals and population splits during the late Paleolithic and Neolithic periods, via Multiple Sequentially Markovian coalescent models and BEAST-based Y-chromosome phylogenies ^21,23,33,63^. These studies also explored fine-scale human population structure and complex migration events. However, the early demographic history of *H. pylori*, such as its origin, initial expansion, bottlenecks, subsequent dispersal, and correlation with human genetic diversity patterns, remains poorly understood because of the underrepresentation of key lineages. To address this, we analyzed the spatial distributions of the major clusters and sublineages identified in our dataset (**Fig. 3a and Extended Data Fig. 5a**). Kriging-based contour maps displayed transparent spatial gradients for each strain. The two highland strains, hspHAEAsiaN and hspHAEAsiaS, had the highest predicted frequencies in the northern and southern Himalayan regions, respectively, which is consistent with their distinct human genetic origins and population structures ^64^. Notably, hspHAEAsiaN also presented elevated frequencies on the northeastern Qinghai–Tibet Plateau and in the upper Yellow River valley, indicating possible ancestral origins and migration routes ^65,66^. In contrast, hspLASEAsia and hspLAEAJapan reached peak frequencies in Southeast Asia and Japan, respectively.

**Fig. 3.**
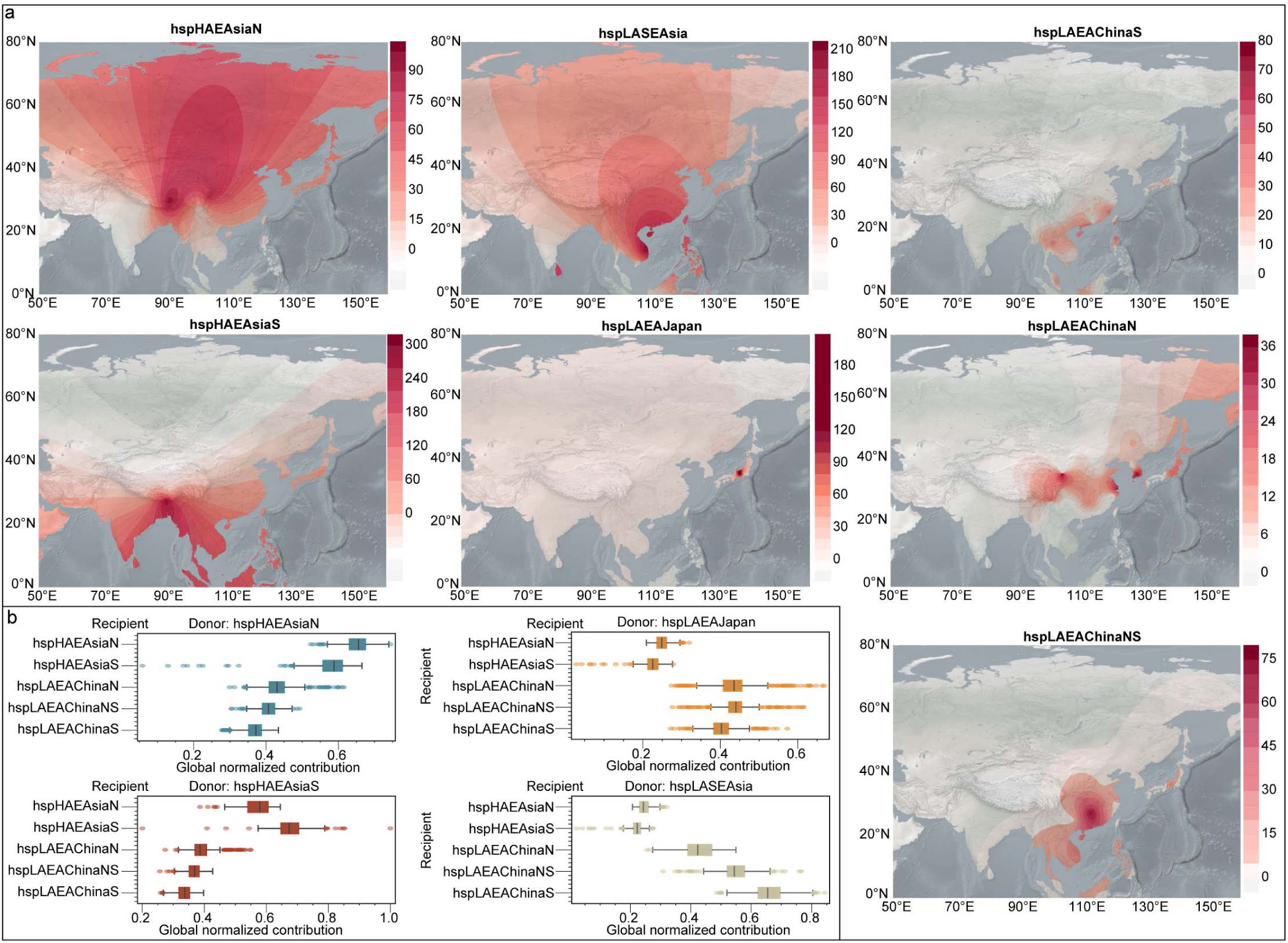
Spatial structure of the hpEAsia lineage and ancestral profile of low-altitude isolates. **a**, Spatial distribution of the seven hpEAsia subtypes. Contour maps depict interpolated frequency surfaces generated with Kriging. **b,** Box-and-whisker plots showing ancestry contributions from selected donors to three recipient groups, including hspLAEAChinaS, hspLAEAChinaN, and the admixed hspLAEAChinaNS. Each panel represents one donor; the x-axis reports globally normalized contributions 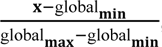, and the y-axis lists recipient categories. Boxes indicate the interquartile range, whiskers span the full data range, and dots denote outliers. Only the donors with the highest contributions are shown; the complete set is provided in **Extended Data Fig. 6**.

Within the hspLAEAChina lineage, the northern sublineages were distributed mainly across the Yellow River Basin, northeastern China, and the Korean Peninsula. The secondary concentration in the Tibetan–Yi Corridor suggests southward migration from northern to southwestern China. The southern lineage was primarily found in the Zhujiang River Basin, along China’s southern coast, and in Taiwan, with additional occurrences on certain Southeast Asian islands, indicating further southward dispersal from southern China into Southeast Asia. In the Yangtze River Basin and Central China, we observed high levels of admixture between the northern and southern sublineages. Correlation analyses between frequency and latitude and longitude for these sublineages revealed that the frequency of hspLAEAChinaN was positively correlated with latitude (R = 0.421, *p* = 0.029), whereas the frequency of hspLAEAChinaS was negatively correlated with latitude (R = -0.415, *p* = 0.044); neither lineage had a significant correlation with longitude **(Extended Data Fig. 5b**). These findings further support distinct genetic differences within the hspLAEAChina lineage. These patterns may reflect human demographic expansions linked to ancient millet farmers in northern China and rice cultivators in southern China ^65,67^, as well as geography-related population substructures inferred from Y-chromosomes and ancient DNA ^21–23,38^. To further explore the genomic origins of the three Chinese lowland sublineages, we used ChromoPainter to infer haplotype sharing with geographically neighboring donor populations (**Fig. 3b; Extended Data Fig. 6**). We observed a north‒south gradient in genetic input: hspLAEAChinaN was enriched for highland-derived haplotypes, whereas hspLAEAChinaS drew mainly from hspLASEAsia. The intermediate group, hspLAEAChinaNS, located in the transitional zone, showed mixed ancestry. Although primarily East Asian, these sublineages also included elements from non-East Asian sources; northern strains showed contributions from hspLAEAJapan, hpNorthAsia, and hpSiberia, whereas southern strains shared features with Southeast Asian lineages ^22,67^. Additionally, these admixture patterns support hypotheses of southward expansion by southern Chinese farmers into Southeast Asia and the northward influence of millet-farming populations on the Qinghai-Xizang Plateau ^65^. Overall, widespread gene flow highlights the significant role of geography in shaping the evolutionary paths of H. pylori in East Asia.

To clarify the evolutionary relationships and phylogenetic history of globally representative *H. pylori* lineages, we performed maximum likelihood (ML) phylogenetic analysis to determine broad clustering patterns (**Fig. 4a and b; Extended Data Fig. 7a**). The clustering of non-Asian lineages matched recent reports by Tourrette *et al*.^16^. Notably, hpAsia2 diverged earlier than other Asian strains did and formed a distinct phylogenetic branch. Most other Asian strains were grouped into a large clade that shared a most recent common ancestor (MRCA) with hpSahul. The complex phylogenetic structure of strains from Siberia and Indigenous American populations, as previously noted, was beyond the scope of this study. Our focus was on East Asian strains, confirming that their clustering reflected geography-related population structure. Three main features emerged: first, multiple sublineages formed separate clades with unique genetic profiles; second, several lineages diverged nearly simultaneously, indicating rapid radiation; and third, despite overall genetic differences, some clades, particularly within hspLAEAChina, included other subgroups. This suggests extensive demographic interactions in East China, with frequent recombination leading to a mosaic evolutionary pattern in certain genomic regions. These findings were supported by an admixture graph generated with TreeMix (**Extended Data Fig. 7b**). The TreeMix analysis of Asian strains revealed that a model with five migration edges (m = 5) explained most of the variance, with additional edges offering minimal improvements (< 0.05%). The topology closely matched the ML tree, with no substantial differences in population classification into major clades. Notably, hpEAsia populations presented short drift branch lengths, indicating limited genetic drift since divergence; this may reflect recent splits, large effective populations, or ongoing gene flow. Migration from the hpAsia2 ecotype into hspHAEAsia was also observed, implying historical genetic exchange between South Asia and the East Asian highlands.

**Fig. 4.**
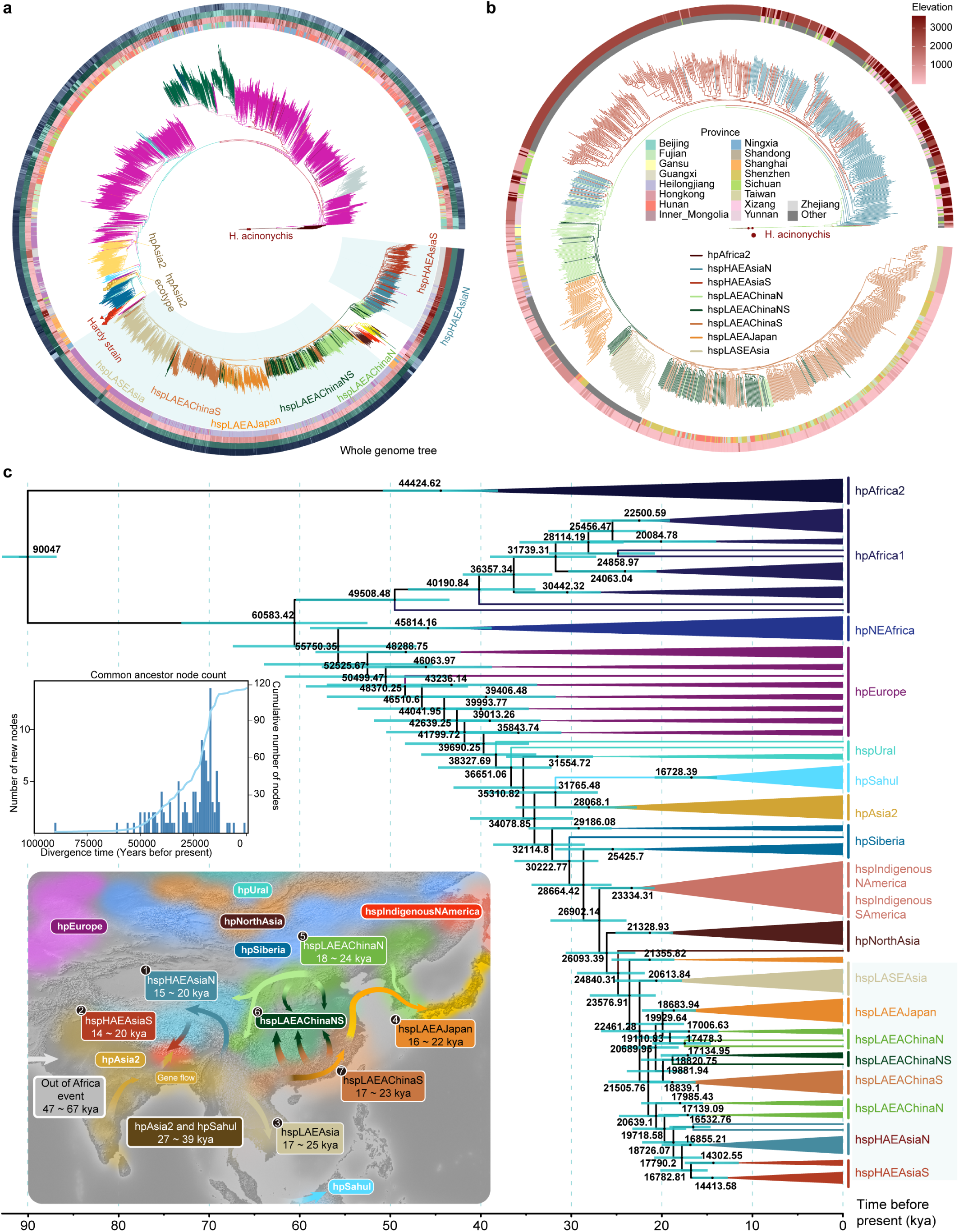
Phylogenetic analysis of *H. pylori* strains. **a**, Maximum-likelihood phylogenetic tree based on whole-genome sequences of the strains analyzed in this study. Branches are colored according to population. *H. acinonychis* serves as the outgroup, as indicated by a red square at the tip of its branch. Hardy strains are marked by red triangle dots at the branch tips, and the hpAsia2 ecotype is marked by a yellow square at the tip of its branch. The concentric circles surrounding each tree represent, from innermost to outermost, the sampling location’s classification by continent, altitude, latitude, and longitude (for details, see **Extended Data Fig. 7a**). **b**, Phylogenetic tree of all hpEAsia strains based on whole-genome sequences. **c**, Time-scaled global phylogeny inferred by BEAST. Individual strains are colored according to subpopulation. At each node, light blue bars indicate the 95% HPD interval, and the mean node age is shown as a numerical label. The upper-left inset depicts the temporal distribution and demographic trajectory of ancestral nodes over successive intervals; the lower-left inset shows inferred migration routes and the timing of *H. pylori* dispersal events.

To estimate divergence times, we reconstructed phylogenies via both ML and Bayesian inference (**Fig. 4c; Extended Data Fig. 8a**). The earliest strains in East Asia seem to be ancient, as both hpAsia2 and hpSahul are found in this region. Bayesian analysis estimated the MRCA of hpAsia2 and hpSahul to date back to 31.8 thousand years ago (kya) (95% highest posterior density [HPD]: 27.1–39.4 kya). Considering their current geographic distributions, these findings support a “southern dispersal” route for early human migration from Africa to the coastal regions of South and Southeast Asia ^68–70^. The MRCA of all hpEAsia strains was dated to approximately 23.6 kya (95% HPD: 21.9–29.4 kya), with the split between hspHAEAsia and hspLAEAsia estimated at 18.8 kya (95% HPD: 16.4–22.2 kya). The ML-based estimates were similar at 29.3 kya and 25.3 kya (**Extended Data Fig. 8b**). These results suggest at least two waves of human migration into East Asia, as shown in the bacterial phylogeny. Notably, the divergence between high- and low-altitude strains had already occurred by the Upper Paleolithic. Historical population dynamics revealed that the effective population size of global *H. pylori* increased in two phases: initially from the late Pleistocene to just before the Last Glacial Maximum (LGM), and subsequently from the LGM to just before the Younger Dryas (**Extended Data Fig. 9**). These expansions imply that climate-driven changes likely influenced human dietary habits and lifestyle shifts, which were later reflected in the *H. pylori* genome.

### Biological adaptive signatures under varying geographic and environmental factors

Having established a strong hierarchical division and detailed population substructure of hpEAsia *H. pylori* populations, we aimed to identify loci and genes with distinctive allele frequency patterns linked to environmental and dietary differences between highland Tibetan and lowland East Asian hosts, as well as adaptive divergence between northern and southern East Asian lineages. Since PC1 and PC2 reflect altitude and latitude differences (**Extended Data Table 3**), we first compared the hspHAEAsia and hspLAEAsia lineages (**Fig. 5a**). This comparison revealed 256 SNPs with *F*_ST_ > 0.6, including 60 SNPs that also surpassed the GWAS significance threshold (−log_10_*p* > 5) (**Fig. 5b; Extended Data Table 4**). The strongest signals overlapped with neuraminyllactose-binding, hemagglutinin-like adhesin, and outer membrane protein loci *alpA*/*hopC*. Additional notable loci with high scores included *frpB4*, *faaA2* (a *vacA*-like cytotoxin), *lysE*, *hopE*, and *hofF*. These genes are associated mainly with surface adhesion and iron acquisition, indicating functional adaptations to high-altitude environments. To investigate latitude-related differentiation, we performed a similar analysis comparing the hspLAEAChinaS and hspLAEAChinaN groups. This analysis revealed that 11 SNPs with *F*_ST_ > 0.6 and 65 SNPs reached GWAS significance (−log_10_*p* > 5) (**Fig. 5b; Extended Data Table 4**). Significant signals include the ribosome recycling factor, the cag-pathogenicity-island component *cagM*, and the outer membrane protein *hofC*. These findings suggest that immune modulation and cell-surface variation may play roles in the north–south divergence of *H. pylori* across East Asia.

**Fig. 5.**
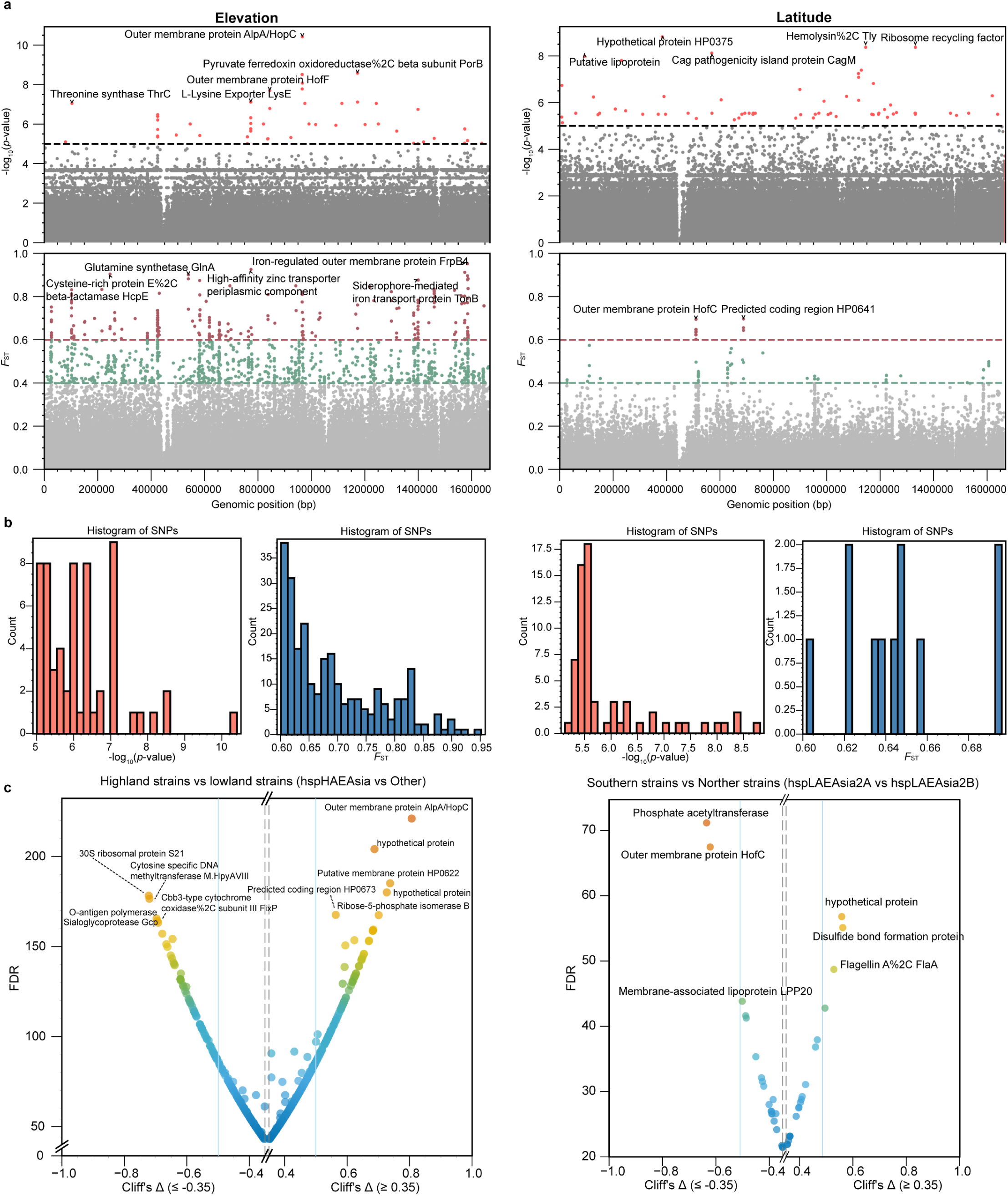
Altitude- and latitude-associated divergent loci in hpEAsia strains. **a**, GWAS and *F*_ST_ scans for variants correlated with altitude and latitude. The GWAS significance is indicated by a horizontal threshold at −log_10_*p* = 5; sites exceeding this threshold are colored red, and those below it are gray. *F*_ST_ thresholds of 0.4 and 0.6 are marked, with SNPs colored gray (< 0.4), green (0.4 ≤ *F*_ST_ < 0.6) or red (**≥** 0.6). **b,** Counts of SNPs surpassing the respective GWAS or *F*_ST_ thresholds. The x-axis denotes −log_10_*p* (GWAS) or *F*_ST_, and the y-axis denotes the number of SNPs. **c,** *dN/dS* analysis per CDS. Only CDSs harboring altitude- or latitude-associated variants are shown; gene products were predicted with Prokka. The x-axis shows Cliff’s Δ, and the y-axis shows the FDR-adjusted −log_10_*p* (see **Methods**). The results with |Cliff’s Δ| > 0.35 are displayed; the ten CDSs with |Cliff’s Δ| > 0.5 are labeled.

To evaluate whether the observed patterns of varied genomic diversity indicate adaptive evolution, we calculated *dN*/*dS* ratios for both the combined coding sequence alignment (**Extended Data Fig. 10**) and individual CDSs (**Fig. 5c**). Among all lineages, hpEAsia presented a higher average *dN*/*dS* ratio than strains from Europe and the Americas did, with hspLAEAJapan exhibiting the highest overall value. Within hspHAEAsia, increased *dN*/*dS* ratios were observed for *alpA*/*hopC*, the putative membrane protein-encoding gene *HP0622*, and *rpiB* (ribose-5-phosphate isomerase), whereas lower ratios were detected for genes encoding the 30S ribosomal protein S21, the DNA methyltransferase M. HpyAVIII, and the O-antigen polymerase (**Fig. 5c**). When the data were stratified by latitude, distinct signals of selection emerged. Genes involved in disulfide bond formation, including *flaA* (flagellin), phosphate acetyltransferase genes, and *hofC*, were targeted differently, indicating divergent selective pressures between northern and southern East Asian strains. To examine the functional basis of these adaptations, we annotated 1,577 coding sequences to identify pathways distinguishing high-altitude from low-altitude strains, as well as northern from southern populations. Despite the use of multiple annotation pipelines, 558 genes lacked Gene Ontology (GO) terms, and 446 lacked Kyoto Encyclopedia of Genes and Genomes (KEGG) annotations, emphasizing the large fraction of *H. pylori* genes that remain uncharacterized (**Extended Data Fig. 11a**). We then overlapped genes with SNPs that passed both the GWAS (−log_10_*p* > 5) and the *F*_ST_ > 0.6 thresholds with those showing |Cliff’s Δ| ≥ 0.35 in the *dN*/*dS* analysis to define a set of candidate targets. All other annotated genes served as the background for subsequent enrichment analyses.

Among highland–lowland high-differentiation adaptive loci (**Extended Data Fig. 11b**), no GO term or KEGG pathway remained significant after applying Benjamini–Hochberg correction (FDR < 0.05), indicating limited statistical power. However, several GO categories showed nominal enrichment (uncorrected *p* < 0.01). Among the top ten GO terms, eight represented biological processes, and two represented molecular functions. The enriched biological processes included siderophore transport (*p* = 1.9 × 10⁻⁴), iron coordination-entity transport (*p* = 6.9 × 10⁻⁴), and iron-ion transport (*p* = 1.0 × 10⁻³), along with nitrogen metabolism, specifically glutamine biosynthetic processes and nitrogen utilization (*p* = 4.9 × 10⁻³). In terms of molecular function, nominal enrichment was observed for energy-transducer activity and phosphomannomutase activity. Although none of these enrichments were significant after multiple testing correction, they aligned with recurrent signals of metal acquisition and surface adhesion identified through GWAS, *F*_ST_, and *dN*/*dS* analyses. In the KEGG database, no pathways achieved significance after FDR correction. Nevertheless, pathways involved in carbon and nitrogen metabolism, two-component signaling systems, and *H. pylori* epithelial cell signaling presented the highest fold-enrichment values, albeit nominal, converging on processes related to metal uptake, nitrogen metabolism, and adhesion. In the analysis of loci with significant north–south differentiation (**Extended Data Fig. 11c**), GO enrichment revealed no terms passing the FDR < 0.05 threshold. However, several categories were nominally enriched (uncorrected *p* < 0.05), mainly related to membrane transport and extracellular structure. These included efflux pump complexes, wide-pore channel activity, porin activity, and efflux transmembrane transporters, as well as flagellar structures such as bacterial-type flagella and cell projections, along with various transporter complexes. After trivial entries (*p* = 1 or fold enrichment = 0) were removed, three pathways were identified: a two-component system (fold enrichment = 25.7, *p* = 1.5 × 10⁻³), flagellar assembly (fold enrichment = 14.3, *p* = 6.9 × 10⁻²), and a bacterial secretion system (fold enrichment = 12.9, *p* = 7.6 × 10⁻²). Although these genes did not reach formal statistical significance, their focus on processes related to transmembrane transport (efflux/porin), motility (flagellum), and signal transduction (two-component systems) matches the genetic signals observed at *cagM*, *hofC*, and genes encoding flagellin and disulfide-bond enzymes. Overall, these results suggest that variation in cell-surface components and environmental sensing mechanisms may influence north–south genomic differences.

We expanded the environmental association analysis to include temperature, solar radiation, and water-vapor pressure, identifying numerous additional SNPs (**Extended Data Fig. 12a**). To reduce collinearity among the environmental variables, we performed PCA on the full environmental matrix (**Extended Data Fig. 12b; Extended Data Table 5**). The first two PCs explained 57.70% and 26.77% of the total variance, respectively. PC1 had positive loadings for temperature (+0.56), water-vapour pressure (+0.47), and solar radiation (+0.46) and negative loadings for latitude (−0.48) and altitude (−0.16), capturing a temperature–humidity–insolation gradient. Higher PC1 scores corresponded to warmer, moister, and more irradiated environments. PC2 was driven primarily by altitude (+0.75) and solar radiation (+0.45), reflecting an altitude–radiation gradient, with higher scores indicating high-altitude, high-irradiance settings. We used PC1 and PC2 as composite environmental predictors in subsequent genome-wide scans (**Extended Data Fig. 12c**). PC1-based GWAS and *F*_ST_ analyses revealed clusters of significant SNPs in genes encoding the flagellar basal-body P-ring protein *flgI*, 5′-methylthioadenosine/S-adenosylhomocysteine nucleosidase, the outer-membrane protein *hopL*, type II citrate synthase *gltA*, and *alpA/hopC*. In contrast, PC2-based scans were dominated by signals in *frpB4*, the GTP-binding protein *obgE*, the vacA-like cytotoxin *faaA2*, a haemagglutinin-like protein, and *vacA*. These findings indicate that both geographic distance and localized environmental pressures jointly shape the population structure and adaptive landscape of *H. pylori* across East Asia. The resulting catalog of candidate SNPs and their genomic contexts provides a critical foundation for future functional studies aimed at elucidating microbial fitness traits, host–pathogen interactions, and disease potential in this globally relevant pathogen.

### Highly differentiated genomic variation shapes distinct clinical outcomes

We first demonstrated that *H. pylori* populations from high and low altitudes exhibit marked genetic stratification across multiple loci. To assess whether altitude-associated variants are linked to clinical traits, we analyzed 106 isolates from Lhasa and 133 isolates from Chengdu (**Extended Data Table 6**). Strain ancestry was inferred via ChromoPainter: genomes assigned to hspHAEAsia were classified as “high altitude”, whereas those assigned to hspLAEAsia were classified as “low altitude". Clinical traits were categorized into two groups (**Extended Data Table 6**): (1) host gastric pathologies and (2) bacterial antibiotic resistance. For each trait, we performed both a GWAS and an *F*_ST_ scan (**Extended Data Fig. 13a, b**), selected loci in the top 5% of each analysis based on empirical thresholds, and then intersected these with the top 5% of loci from the high-versus-low-altitude differentiation scan. We then filtered out variants implicated in both altitude-related differentiation and clinical outcomes and compared their allele frequencies between the high-altitude (hspHAEAsia) and low-altitude (hspLAEAsia) groups, as well as among individual strain populations (**Fig. 6a; Extended Data Fig. 13c**). A two-sample t-test confirmed highly significant differences in allele frequency between hspHAEAsia and hspLAEAsia (*p* < 0.0001), demonstrating that these loci were indeed differentiated by altitude. Moreover, both the hspHAEAsiaN and hspHAEAsiaS subpopulations presented higher allele frequencies than did all the other groups; overall, the hspLAEAChina population presented higher frequencies than hspLASEAsia and hspLAEAJapan did. We then constructed contingency tables stratified by altitude and phenotype, applying Pearson’s χ² test or Fisher’s exact test as appropriate (see **Materials and Methods**). Odds ratios (ORs) were derived from the same contingency tables, and SNPs with *p* < 0.05 were considered significant. In total, 489 SNPs met this criterion; of these, 464 could be annotated by Prokka and mapped to 175 distinct genes (**Fig. 6b; Extended Data Table 7**). High ORs indicated strong positive associations between minor alleles and clinical traits. Notable examples included position 1,299,728 in the d-lactate dehydrogenase gene, associated with tetracycline resistance (OR = 136.5); position 1,586,955 in *frpB4*, associated with peptic ulcers (OR = 132); and position 1,240,595 in *gluP*, associated with intestinal metaplasia (OR = 101.7) (**Fig. 6b**). These SNPs, which vary in frequency between high- and low-altitude populations and correlate with distinct clinical outcomes, represent candidate risk alleles for future functional validation.

**Fig. 6.**
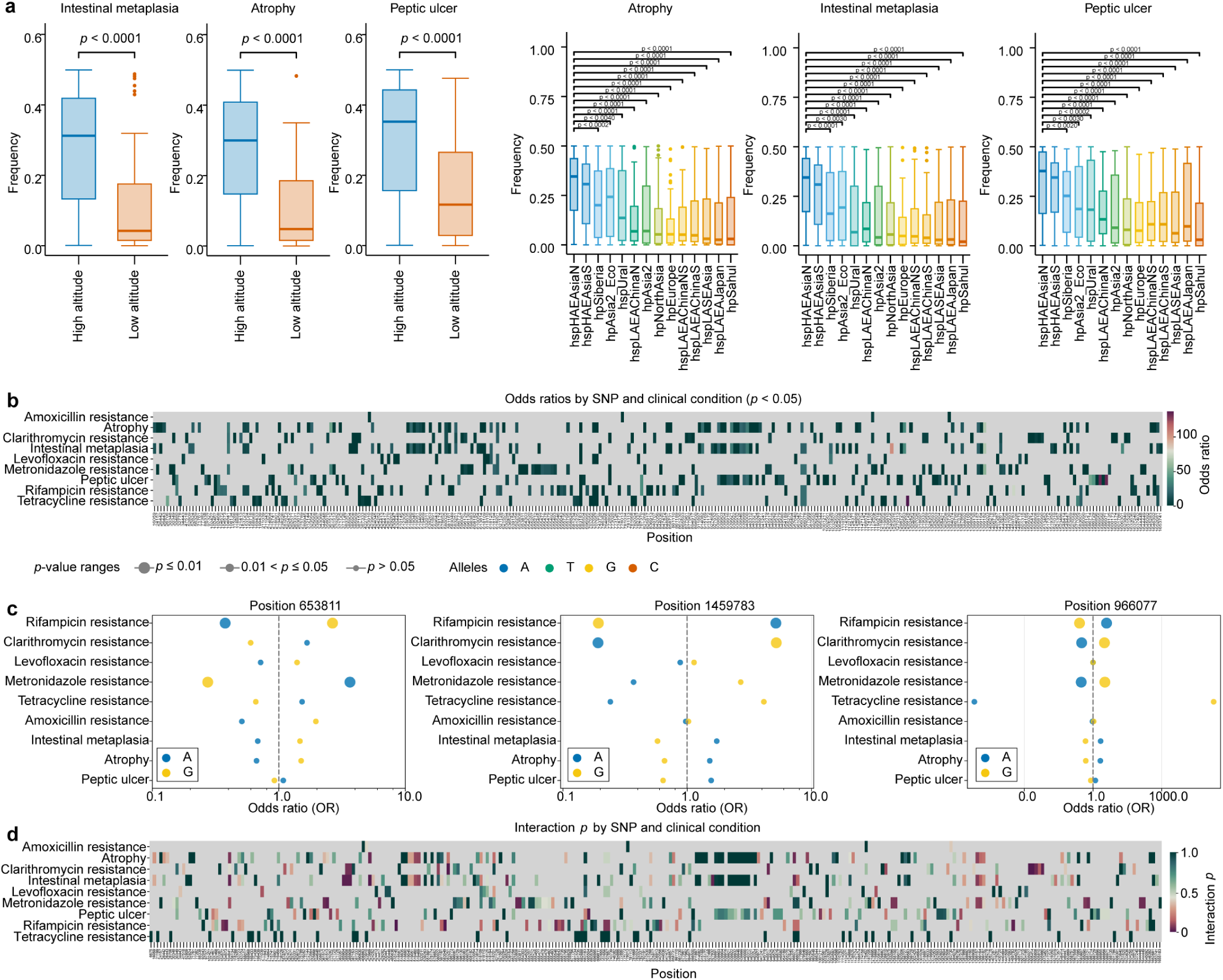
Landscape of SNP variants associated with host clinical and antibiotic resistance phenotypes. **a**, Boxplots of allele frequency distributions of candidate loci across populations stratified by host clinical phenotype (peptic ulcer, intestinal metaplasia, gastric atrophy). Candidate loci are defined as the top 5% of SNPs from GWAS and *F*_ST_ analyses and intersect with the top 5% of loci from a high-versus-low-altitude differentiation scan; allele frequencies were computed for each locus within each population, and the statistical significance was tested using a two-tailed t-test.. The distributions of loci associated with bacterial antibiotic resistance phenotypes are shown in **Extended Data Fig. 13c. b**, Odds ratios (ORs) from χ² or Fisher’s exact tests (*p* < 0.05); color intensity represents OR magnitude on a custom linear scale, the color bar indicates OR values, and gray denotes non-significant associations. **c,** Bubble plot of ORs for pleiotropic SNPs. The x-axis shows ORs (log scale); the y-axis lists phenotypes. The bubble color indicates the allele (A or G), and the bubble size corresponds to the Bonferroni-adjusted *p*-value category (*p* ≤ 0.01; 0.01 < *p* ≤ 0.05; *p* > 0.05). The dashed vertical line indicates OR = 1. ORs and *p*-values were obtained from logistic regression with Wald tests and Bonferroni correction; the remaining loci are shown in **Extended Data Fig. 14. d**, Heatmap of interaction *p*-values for SNP–phenotype associations; color intensity reflects *p*-value magnitude, and gray indicates non-significance.

Several loci were associated with more than one clinical phenotype. To investigate these pleiotropic effects, we applied logistic regression models to each SNP across multiple traits and identified 322 SNPs associated with at least two clinical phenotypes. Of these, 21 exhibited significant SNP–phenotype interaction effects (unadjusted *p* < 0.05), and following Bonferroni correction, 15 remained significant (Bonferroni-adjusted *p* < 0.05) (**Extended Data Table 8; Extended Data Fig. 14**), 14 of which mapped to nine distinct gene regions. Notably, SNP 653,811 in *faaA2* (putative VacA-like toxin) was significantly associated with resistance to both rifampicin and metronidazole (*p* < 0.05 for each) (**Fig. 6c**). Similarly, the A allele at position 1,459,783 conferred increased rifampicin resistance (OR = 5.18, *p* < 0.05), whereas the G allele at the same site was linked to clarithromycin resistance (OR = 5.22, *p* < 0.05). SNP 966,077 in *alpA/hopC* correlated with resistance to rifampicin, clarithromycin, and metronidazole (*p* < 0.05 for all). These findings identify a set of pleiotropic variants whose mechanistic underpinnings warrant further investigation.

To assess whether the clinical impact of specific variants is influenced by the genomic background shaped by altitude, we implemented genotype‒altitude interaction models for each SNP, and we retained 20 SNPs with interactions *p* < 0.05, 19 of which mapped to 12 distinct gene regions (**Fig. 6d; Extended Data Table 9**). These SNPs indicated that the clinical consequences of identical mutations varied between high- and low-altitude strains. For rifampicin resistance, mutations at position 510,145 (G>T) in *hofC* (OR = 14.3) and at position 583,049 (G>T) in *cagA* (OR = 11.7) significantly increased rifampicin resistance in high-altitude isolates. Similarly, a variant at position 385,904 (A>G) in *dsbC* was strongly associated with an elevated risk of intestinal metaplasia in high-altitude hosts (OR = 11.7). Together, these altitude-dependent variants highlight promising candidates for functional validation and offer potential biomarkers for precision diagnostics, individualized treatment, and region-specific clinical management strategies.

### Integrated database and query platform

To address the fragmentation and inconsistent updates among existing resources, we developed a unified, web-based database and query platform that enables rapid access to genome-wide SNP variation data in *H. pylori* (**Fig. 7a-g**). Users can input a genomic coordinate corresponding to an *H. pylori* SNP site, prompting the system to retrieve variation data across 7,544 whole-genome sequences. The platform then computes the count and frequency of each allele, accounting for the possibility of multiple alleles at a single locus, under six classification schemes: Subpopulation: 26 groups defined in this study, including six newly reclassified lineages (hspLASEAsia, hspLAEAChinaS, hspLAEAChinaNS, hspLAEAChinaN, hspHAEAsiaN, and hspHAEAsiaS) derived from the former hpEAsia, one hpAsia2 ecotype subset, and 19 lineages adopted from previous studies. Geographical bubble map, where bubble size reflects allele count and color represents allele type at each location. An animated density map illustrates allele frequency (allele count divided by total count at each site) across geographic coordinates. There is also information about the main population, country, and continent. By providing immediate visualization of the global allele distribution at any locus, the platform enables researchers to identify lineage- or region-specific SNPs, thereby accelerating the generation of hypotheses, experimental design, and development of targeted public health strategies. The platform also incorporates an *H. pylori* reference panel module for genotype imputation (**Fig. 7h**). Users can upload draft assemblies, after which the cloud server imputes missing genotypes and returns a complete whole-genome file. To support local analyses and reduce the server load, the reference panel itself is available for download and offline imputation.

**Fig. 7.**
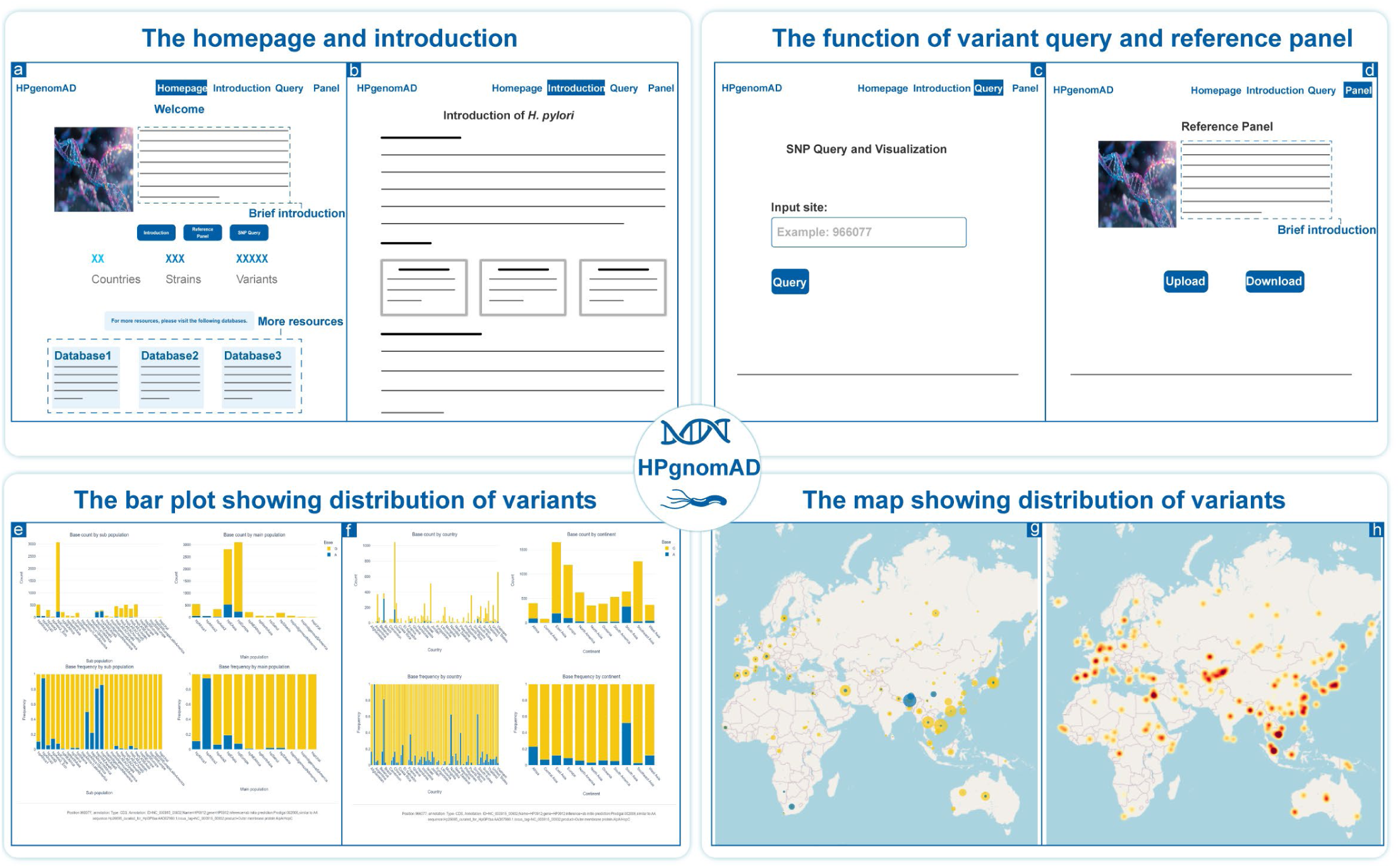
Web interface of HPgnomAD. **a**, Landing page providing an at-a-glance summary of the underlying dataset. **b,** Introductory module describing *H. pylori* biology and epidemiology. **c,** Query panel for entering a SNP position relative to strain 26695. **d,** Interactive results stratified by population and subpopulation, showing both absolute counts and allele frequencies. **e,** Interactive results stratified by country and continent, likewise displaying counts and frequencies. **f,** Dynamic map depicting the geographical distribution of allele counts at the queried locus. **g,** Dynamic map depicting the geographical distribution of allele frequencies at the same locus. **h,** Haplotype imputation reference panel and imputation function: users can upload a partial sequence file or download the reference panel for imputation.

## Discussion

### Fine-scale population substructure and deep demographic history of *H. pylori*

*H. pylori* colonizes the human stomach almost exclusively, with infection recognized as the principal etiological driver of atrophic gastritis and a major risk factor for peptic ulcer disease and gastric carcinoma. Genomic analysis of *H. pylori* offers a dual lens, illuminating human origins, migrations, and evolutionary history while clarifying host–pathogen co-evolution. These insights provide a robust foundation for disease prevention, precision therapy, and personalized medicine. Epidemiological data indicate that gastric inflammation severity and gastric cancer risk are markedly greater in Asia, particularly East Asia, than in Europe and North America, a disparity likely driven by more virulent *cagA* and *vacA* genotypes ^71–73^. This underscores the urgency of comprehensively characterizing East Asian *H. pylori* strains. Earlier genomic investigations, constrained by available technologies, relied heavily on MLST, pulsed-field gel electrophoresis, random amplified polymorphic DNA, and amplified fragment length polymorphism, with MLST as the dominant method. MLST, which targets seven housekeeping genes, enables the classification of seven modern and six ancestral *H. pylori* populations ^74^, but its limited genomic scope restricts resolution. The advent of high-throughput WGS supplanted MLST, with the first *H. pylori* genome sequenced by Tomb *et al*. in 1997 ^9^, ushering in fine-scale reconstructions of global population structure and migratory history ^15,16,29,30,51,58^. Despite East Asia’s vast geography, demographic complexity, and host genetic diversity, most studies have continued to treat its *H. pylori* strains as a single hpEAsia (or hspEAsia) population ^15,16^. Although some studies have identified regional sublineages and proposed location-based nomenclatures ^29,30,75^, these efforts have been limited by regional sampling, incomplete temporal resolution, and the continued use of MLST-based definitions, which obscure whole-genome diversity and microvariation. The Qinghai–Tibet Plateau, a high-altitude, hypoxic region with a distinct human population history, remains a major gap, with scarce isolates hindering evolutionary reconstruction in this extreme environment. To address these gaps, we assembled the largest *H. pylori* genomic dataset to date, integrating 7,544 publicly available genomes. Using fineSTRUCTURE, PCA, and ADMIXTURE, we present the first genome-wide hierarchical subdivision of hpEAsia, revealing pronounced differentiation between high- and low-altitude communities and along a north–south latitudinal gradient.

Our expanded dataset uncovers a far more complex hpEAsia substructure than previously recognized. To avoid geographic nomenclature that obscures relationships, we adopt an alphanumeric scheme. At the broadest scale, hpEAsia segregates into two altitude-defined lineages: hspHAEAsia (high-altitude) and hspLAEAsia (low-altitude). hspHAEAsia is confined to the Qinghai– Xizang Plateau and adjacent regions, comprising hspHAEAsiaN (Tibet, China) and hspHAEAsiaS (Bhutan). hspLAEAsia includes three main sublineages—hspLASEAsia (predominant in Southeast Asia), hspLAEAChina (widespread in mainland China), and hspLAEAJapan (largely restricted to Japan). Within hspLAEAChina, ADMIXTURE, fineSTRUCTURE, and phylogeny revealed additional fine-scale substructures including hspLAEAChinaS, hspLAEAChinaN, and an admixed hspLAEAChinaNS. Regression analyses confirmed that both altitude and latitude significantly influence genomic variation, with altitude exerting a stronger effect. While the hspHAEAsia–hspLAEAsia split remains the deepest division, a north‒south gradient persists within hspLAEAChina, with transitional populations reflecting continuous gene flow among mainland Chinese hosts. ChromoPainter analysis revealed that each hspLAEAChina subtype derives more genomic segments from nearby strains than from distant strains, mirroring historical human migration and bacterial exchange. In addition to these subdivisions, we identified additional genomic divergence and a novel ecotype within hpAsia2. Tourrette *et al*. defined an ecotype as a bacterial population that maintains a single recombining gene pool but exhibits speciation-level divergence in specific genomic regions ^16^. In the study of hspIndigenousSAmerica and hspIndigenousNAmerica, they reported highly recombining yet strongly differentiated genomic segments, challenging classical species-formation models. Our findings demonstrate that *H. pylori* ecotypes are widespread, expanding the ecological and evolutionary framework for ecotype diversification.

TMRCA estimates from ML and Bayesian analyses date the split between hpAfrica2 and all other lineages to ∼90.0 kya (95% HPD: 86.9–92.9 kya). hpNEAfrica and non-African strains diverged at 55.8 kya (95% HPD: 52.5–73.1 kya), which aligns with previous *H. pylori* ^76^, human Y-chromosome ^77^ and mtDNA ^78^ estimates. The earliest Asian lineage, hpAsia2, diverged from hpSahul ∼31.8 kya (95% HPD: 27.1–39.4 kya), implying modern human expansion into South Asia and the Sahul by that time. hpEAsia’s TMRCA is ∼23.6 kya (95% HPD: 21.9–29.4 kya), with the hspHAEAsia–hspLAEAsia split at ∼18.8 kya (95% HPD: 16.4–22.2 kya), firmly in the Paleolithic. Archaeological ^79,80^ and genetic ^81–83^ evidence support high-altitude habitation on the Qinghai–Xizang Plateau by ∼18.8 kya. Ancient DNA ^23,84^ further documented a north–south East Asian split by ≥ 19 kya, matching the diversification of hspLAEAChina. Together, pathogen and host evidence converge on both altitudinal and latitudinal differentiation in East Asian human-*H. pylori* histories. Reconstructing *H. pylori* evolutionary relationships, however, requires careful treatment of three challenges: pervasive recombination, extreme genetic diversity, and molecular clock uncertainty ^44,85,86^. First, *H. pylori*’s high recombination rate introduces more variation than de novo mutations do. While tools such as ClonalFrameML can remove recombinant segments for divergence dating, this often erases recent demographic signals, producing artificially flat BSPs ^87^. Second, *H. pylori*’s small yet highly diverse genome (∼1.6 Mb) yields a nonlinear increase in segregating sites with larger sample sizes, creating computational bottlenecks for Bayesian algorithms. Third, mutation rate estimates remain incomplete, with microevolutionary rates exceeding macroevolutionary rates because of purifying selection ^44,88^. Robust molecular dating demands calibration against well-dated historical or geological events, with ancient DNA providing key opportunities ^89^, as in human mtDNA ^90–92^ and *Mycobacterium tuberculosis* ^93^ studies. Future work should focus on developing frameworks that integrate recombining and nonrecombining genome regions to infer evolutionary history and gene flow jointly, designing phylogenetic algorithms optimized for large-scale WGS analyses, and refining molecular-clock estimates through calibration with ancient *H. pylori* genomes.

### Biological adaptive signatures and clinical significance

Previous studies reported a greater risk of *H. pylori* infection in Tibetan populations than in Han populations ^94^. An MLST-based analysis attributed this susceptibility to a Western-type (hpEurope) *cagA* allele in Tibetan isolates ^95^. In contrast, our whole-genome analyses revealed that plateau strains belong almost exclusively to hspHAEAsiaN and showed no clustering with hpEurope in either genome-wide or core-CDS phylogenies, underscoring the necessity of WGS for accurate strain classification. To pinpoint the genomic features driving high-altitude versus low-altitude differentiation, we performed a GWAS and calculated the *F*_ST_, identifying numerous highly differentiated candidate loci likely contributing to increased virulence in plateau strains. *dN*/*dS* analysis highlighted a set of genes encoding hemagglutinin-like adhesin, the outer-membrane protein *alpA*/*hopC*, *frpB4*, and several iron-uptake system components. GO enrichment indicated nominal (uncorrected *p* < 0.01) overrepresentation of iron-transport terms, including siderophore transmembrane transport (*p* = 1.9 × 10^−4^), iron–ligand transmembrane transport (*p* = 6.9 × 10^−4^), and iron ion transport (*p* = 1.0 × 10^−3^), as well as nitrogen metabolism categories. Although these signals did not survive multiple testing correction, they align with the GWAS, *F*_ST_, and *dN*/*dS* results, suggesting that plateau strains adapt to the low-iron gastric environment by optimizing iron acquisition and adhesion, enhancing colonization and pathogenicity.

North–south genetic differentiation was relatively modest. The most differentiated SNPs in these contrasts mapped to genes involved in cell-surface architecture and signal transduction, such as *cagM*, *hofC*, flagellin, and disulfide bond-forming enzymes, indicating a subtler latitudinal signature of adaptation. We caution against attributing all differentiation solely to altitude or latitude, as correlated environmental factors may also impose selective pressures. Environmental principal component analysis decomposed multiple climatic variables into two components, implicating composite gradients of temperature–humidity–radiation and altitude– radiation as major drivers of *H. pylori* genomic adaptation. These findings lay the groundwork for future functional validation of adaptive evolution in this pathogen. To assess potential clinical impacts, we compiled phenotypic data on antibiotic resistance (tetracycline, rifampin, metronidazole, clarithromycin, etc.) and gastric disease manifestations (peptic ulceration, intestinal metaplasia, etc.). GWAS and *F*_ST_ analyses identified SNPs linked to divergent clinical outcomes, addressing three questions: whether genomic variants distinguishing high- and low-altitude strains are associated with host pathology or drug resistance; whether individual SNPs show pleiotropic effects; and whether identical mutations produce distinct clinical consequences in plateau versus lowland contexts. Multiple candidate SNPs have emerged, with several conferring stronger pathogenic or resistance phenotypes in high-altitude isolates—evidence for genotype–environment interactions shaping clinical presentation. These variants offer promising biomarkers for precision diagnostics and individualized therapy, enabling targeted interventions across ecological contexts. Recognizing the critical role of WGS in resolving population structure and mapping pathogenic loci, we aggregated genome-wide variant calls from 7,544 isolates into an online query platform. This resource allows users to retrieve allele counts and frequency distributions for any SNP across strains and regions, which is supported by interactive visualizations. By integrating large-scale genomic data with population-genomic analyses, we provide new perspectives and tools for elucidating *H. pylori* evolution, environmental adaptation, and clinical diversity, advancing both basic understanding and precision public health strategies.

## Materials and methods

### Sequence collection and quality control

A total of 8,010 *H. pylori* sequences were retrieved from publicly available databases, including NCBI (https://www.ncbi.nlm.nih.gov/), EnteroBase (https://enterobase.warwick.ac.uk/), PubMLST (https://pubmlst.org/organisms/helicobacter-pylori/), and Datadryad (https://datadryad.org/stash). After quality control, 7,544 high-quality sequences (including two H. acinonychis sequences and 7,542 *H. pylori sequences*) were retained. We generated a new *H. pylori* genomes with detailed clinical phenotypes from 106 individuals in Lhasa, Tibet, and 89 individuals in Chengdu, Sichuan, using the Illumina MiSeq platform in this database. To ensure the integrity of downstream variant discovery and annotation, the reference genome was preprocessed by creating a genome index with Samtools and a sequence dictionary with Picard, providing the necessary metadata for variant analysis. Initial variant filtering employed GATK (v4.2.6.1) VariantFiltration with the following thresholds: QUAL < 20, QD < 1, MQ < 30, FS > 100, and SOR > 10 ^96^. Variants flagged as low quality were subsequently removed using VCFtools (v0.1.16). To further maximize reliability, we applied stringent criteria, retaining only variants with a missing rate ≤ 5%, minimum read depth ≥ 3, MAF ≥ 5%, average sequencing depth ≥ 20, and QUAL ≥ 30. Low-quality genomes were excluded on the basis of the following criteria: assembly fragmentation exceeding 500 contigs and coverage of the reference genome (*H. pylori* strain 26695, NC_000915.1) below 70% using BWA v.0.7.13 ^97^, as described by Tourrette *et al*. ^16^. Genome annotation for each sequence was performed via Prokka v1.14.6 (option: --genus Helicobacter, --species pylori, --kingdom Bacteria, --evalue 10^-24^)^98^.

### Core genome variant calling

The nucmer module in MUMmer v4 software was employed for SNP calling ^99^. Specifically, each genome sequence in the dataset was aligned against the reference genome of *H. pylori* (strain 26695, NC_000915.1). SNPs were identified via the show-snps and show-coords functions with the parameters (-ClrT and -rclT). Delta files and coord files were converted into VCF format as described previously ^16^. BCFtools v.1.14 was used to merge files, summarize variant information, and extract specific variants ^100,101^. In a total of 7,544 *H. pylori* genomes, we identified 1,864,491 SNPs and 497,548 InDels. After a minimum allele frequency threshold of 1% was applied, 829,724 core SNPs and 271,723 InDels were retained. To simplify the alignment, we applied snp-sites v2.5.2 to remove monomorphic positions, retaining only polymorphic sites ^102^. GNU parallel software was used to run multiple serial command line programs in parallel ^103^. The visualizations were generated via the Tidyplots package ^104^.

### Haplotype imputation reference panel construction

For HPgnomAD, raw variants were first processed with BCFtools by removing SNPs located within 3 bp upstream or downstream of indels ^100,101^. Uncertain genotypes were set to missing, and all missing calls were filled with the reference allele to generate a standardized haploid haplotype reference. Non-polymorphic sites (allele count = 0) were then filtered out via BCFtools view. Each sample set was subsequently phased and self-imputed with Beagle 5.0 ^105^. To evaluate imputation performance, we randomly selected 20 individuals per population as validation samples. Within these samples, genotypes were masked at different coverage levels to simulate low-information scenarios, and imputation was performed separately with each panel via Beagle 5.0. Imputation quality was assessed for each SNP by three complementary metrics: (1) dosage correlation squared (DR^2^), as estimated by Beagle on the basis of the ratio of imputed dosage variance to the theoretical variance under known genotypes; (2) empirical dosage R^2^ (R^2^), calculated as the square of the Pearson correlation between imputed and true dosages across all validation samples; (3) the Pearson correlation (r) between the mean dosage before and after imputation; and (4) genotype concordance, defined as the proportion of best guess imputed genotypes that exactly match the true genotypes. By combining these metrics and stratifying by masking level, we quantified the panel’s accuracy and robustness under low coverage or low-density conditions.

### Strain assignment

According to previous studies ^12–14,16,17^, 285 strains were consistently assigned to one of 19 *H. pylori* populations/subpopulations. These 285 strains were previously examined and confirmed to represent their respective populations or subpopulations. To optimize computational efficiency, we randomly selected ten strains from each population, retaining all available strains when fewer than ten existed (**Extended Data Table 1**). We then assembled a dataset of 791 strains and applied fineSTRUCTURE ^106^ to validate both their individual assignments and population assignments. To dissect the genetic substructure of the hpEAsia lineage, we iteratively removed non-hpEAsia strains and repeated the fineSTRUCTURE analyses (**Extended Data Fig. 2**). The fineSTRUCTURE analysis was carried out for 200,000 Markov chain Monte Carlo iterations, including a burn-in of 50,000 iterations, based on core genome SNPs. After verifying the reassignment of each strain, these strains were used as a donor panel to “paint” each sample of remaining hpEAsia via ChromoPainter v2 ^106^. Each genome was then assigned to a population on the basis that the population contributed the largest proportion of its painted genomic segments.

To objectively evaluate the difference in contributions between the hspLAEAChinaS and hspLAEAChinaN components derived from the ChromoPainter analysis, we constructed a differential index defined as follows: 𝛿 = hspLAEAChinaS − hspLAEAChinaN. Statistical analysis of the 𝛿 values from 1443 samples yielded a standard deviation of σ ≈ 249. To distinguish samples with minimal contribution differences, we adopted a relative thresholding approach by setting the threshold as k × σ. In this study, we selected k = 0.5, corresponding to a threshold of approximately 125. Therefore, samples satisfying |𝛿| ≤ 0.5 × σ were considered to exhibit negligible differences between hspLAEAChinaS and hspLAEAChinaN contributions and were subsequently classified into a hybrid group, denoted hspLAEAChinaNS (**Extended Data Fig. 7**). Finally, we delineated 26 populations and subpopulations from 7,544 genomes (**Extended Data Table 1**). We subdivided hpEAsia into seven subtypes, including hspHAEAsiaN, hspHAEAsiaS, hspLAEAChinaN, hspLAEAChinaNS, hspLAEAChinaS, hspLAEAJapan and hspLASEAsia, and identified a distinct ecotype within hpAsia2, designated the hpAsia2 ecotype.

### Principal component analysis

The merged VCF format file was processed via PLINK v1.9 to remove linkage disequilibrium according to the following criteria: window size of 50 base pairs, step size of 10 variants, and an r^2^ threshold of 0.1 ^107^. Visualization was performed via Python v3.12.2, which employs the seaborn and Matplotlib packages in Python ^108–110^.

### Uniform Manifold Approximation and Projection (UMAP)

To investigate the genetic structure of hspHAEAsia and hspLAEAsia, we performed UMAP dimensionality reduction in Python v3.12.2 with the umap-learn package (v0.5.3). The parameters were set as follows: n_neighbors = 10, min_dist = 0.1, n_components = 2, metric = Euclidean, init = spectral, and random_state = 42 ^111^.

### Spatial geographic distribution

The spatial distribution of each strain across geographic Asia was visualized via Surfer^®^ from Golden Software, LLC (www.goldensoftware.com). Contour maps depicting geographic frequency patterns were generated via the Kriging interpolation algorithm implemented in Surfer. Data derived from non-population-based samples were excluded from the analysis to ensure representativeness.

### Genome structure comparison

To compare genome architecture, we employed dot plots to visualize substantial structural variations, such as inversions, gaps, repeats, and gene cluster rearrangements. Dot plots were generated with Gepard (v. 1.40) to compare genomic similarity and divergence between two distinct hpAsia2 lineages within the phylogeny ^112^. Specifically, we carried out pairwise comparisons of the hpAsia2 ecotype versus hpAsia2, hpAsia2 ecotype versus hpAsia2 ecotype, hpAsia2 versus hpAsia2, and hpAsia2 ecotype versus *H. cetorium* (NCBI accession CP003479.1). The alignments and plots were constructed with Gepard’s internal DNA substitution matrix (edna.mat). A lower color threshold of 10 was chosen to suppress background noise and emphasize regions of interest. Both the window size and word length remained at their default settings (0 and 10, respectively).

### Phylogenetic analysis

We conducted phylogenetic analyses of the whole genome and the core coding sequence (CDS) regions. The aligned sequences were then used to construct phylogenetic trees via VeryFastTree v4.0.4 with the -nt option ^113^. The resulting tree files are visualized and refined via the ggtree and ggtreeExtra packages ^114,115^. For each tree, *H. acinonychis* served as an outgroup. To estimate strain divergence times, we applied two established methods ^15,30^, using a stratified random subset of 100 samples from the dataset to increase computational efficiency. In the first approach, we reconstructed a phylogeny via IQ-TREE v3 (options: -m MF, -T AUTO) and Model Finder under the Bayesian information criterion to select the GTR+G substitution model ^116^. We then applied ClonalFrameML v1.12 to these whole-genome sequences under default settings to account for recombination ^117^. Node ages were estimated in PAML v4.10.8 (model = 7, Mgene = 0, ncatG = 4) ^118^ by assuming a uniform mutation rate of 1.38×10^-5^ substitutions per site per year ^44^. In the second approach, recombinant regions were first masked via ClonalFrameML v1.12. The filtered alignment was then analyzed in BEAST2 v2.5.0 ^119^, running 4×10^7^ MCMC iterations under a GTR substitution model with a relaxed lognormal clock and the same mutation rate of 1.38×10^-5^ per site per year.

### Admixture graphs

ADMIXTURE analysis was conducted via ADMIXTURE software ^120^. To optimize computational efficiency, we first subsampled global strains by randomly selecting 20 isolates from each non-hpEAsia population or subpopulation (retaining all available isolates when fewer than 20 were present) and 50 isolates per hpEAsia population. We then evaluated K values from 10 to 20 via eight computational threads, ten-fold cross-validation and a fixed random seed (12345) to ensure reproducibility. Admixture graphs among the different populations were constructed via TreeMix v.1.12 ^121^. The analysis was performed with migration edges ranging from 0 to 15, with 10 replicates for each specified number of edges. The population hpAfrica2 was designated as the outgroup. The optimal number of migration edges was determined as the smallest value that accounted for 99.8% of the explained variance.

### The calculation of nonsynonymous and synonymous substitutions

Nonsynonymous substitutions refer to nucleotide changes that result in amino acid changes, whereas synonymous substitutions are those that do not alter the amino acid sequence. Both substitutions and their ratios were determined through pairwise comparisons of each strain’s genome coding sequences against the hpAfrica2 strain via PAML v4.9 ^118^. The calculations were performed via the Nei‒ Gojobori (1986) method ^122^ and the Yang‒Nielsen (2000) method ^123^.

To pinpoint CDS regions under natural selection in *H. pylori*, we computed *dN/dS* for all 1,577 CDS regions annotated by the reference strain 26695 (NC_000915.1) against the hpAfrica2 strain via the Nei–Gojobori framework. Differences in selective pressure were then assessed via two complementary strategies: (1) the strain of interest was treated as one group, and all remaining strains constituted the second group; (2) two arbitrarily chosen strain sets were compared directly. Statistical tests were restricted to CDS regions represented by ≥ 5 observations in both groups. The distributions of *dN/dS* were contrasted with a two-sided Mann–Whitney U test, yielding U statistics and raw *P* values. The effect size was expressed as Cliff’s Δ, which was calculated as 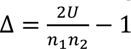, where n_1_and n_2_ denote the sample sizes of the two groups. Multiple testing correction was performed with the Benjamini–Hochberg procedure. CDS regions meeting FDR < 0.05 and |Δ| ≥ 0.33 were deemed to exhibit significant selective differences.

### GWAS and *F*_ST_ analysis

The environmental factors, external conditions influencing organismal development, behavior, and survival, were quantified as follows. Geographic coordinates (latitude and longitude) were first determined from existing location data. The environmental variables for each site were subsequently retrieved from the WorldClim database ^124^, including altitude (m), temperature (°C), solar radiation (kJ m^−2^day^−1^), and water vapor pressure (kPa). Treating environmental factors as quantitative traits, we conducted a GWAS to pinpoint biallelic core SNPs associated with each trait. A total of 250 samples were randomly selected for each trait via a binning-based sampling strategy to ensure uniform coverage of the range of the trait. In brief, trait values were sorted and evenly partitioned into 10 bins, and samples were randomly drawn from each bin. GWAS was conducted via the lin_loc function from the R packages bug (v.0.0.0.9000) and GEMMA (v.0.93) ^125^, employing a significance threshold of -log_10_(*p*-value) = 5. The per-site *F*_ST_ values were computed with the R package PopGenome to reinforce the GWAS findings ^126^. The GWAS samples were dichotomized on the basis of trait values for each trait, and this binary classification was used as the basis for the *F*_ST_ analysis. For the *F*_ST_ comparisons, the isolates were dichotomized into “high” and “low” groups by selecting the top 125 and bottom 125 samples for each continuous variable. To identify loci that jointly underlie high-altitude adaptation and clinically relevant phenotypes, we analyzed 239 *H. pylori* isolates with matched clinical metadata from Chengdu and Lhasa (**Extended Data Table 6**) via the same pipeline. Four metrics were considered: altitude-based GWAS scores, altitude *F*_ST_, phenotype-based GWAS scores and phenotype *F*_ST_. SNPs exceeding the 95th percentile for all four metrics were retained as candidates. Genotypes at each candidate SNP were extracted to construct contingency tables stratified by allele and subgroup (high-altitude cases, high-altitude controls, low-altitude cases and low-altitude controls). Biallelic tables with all expected cell counts ≥ 5 (and with fewer than 20 % of cells < 5) were analyzed by Pearson’s χ² test; otherwise, Fisher’s exact test was applied. ORs comparing high-versus low-altitude allele frequencies were calculated from the corresponding allele contingency tables via Python’s statsmodels according to 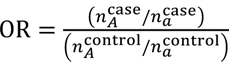, where and denote the counts of alleles A and a, respectively.

To determine whether the effect of genotype on the clinical phenotype differs between high- and low-altitude environments, we fitted, for each SNP, a logistic regression model of the form (phenotype ∼ altitude + genotype + altitude × genotype), extracted the *p* value for the interaction term, and reported the ten SNPs with the most significant genotype–altitude interactions. Finally, to assess whether a given SNP exerts effects across multiple clinical phenotypes, we first assembled an allele–phenotype matrix for each binary outcome. We then fitted separate logistic regression models of the form 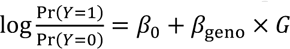, where 𝑌 denotes the binary clinical phenotype and where 𝐺 is the genotype is additively coded. Models were estimated in Python via the statsmodels library, employing the default Newton algorithm until convergence. ORs were computed as exp (𝛽_geno_). To correct for multiple testing across 𝑚 SNPs and 𝑘 phenotypes, we applied a Bonferroni adjustment, setting the significance threshold at 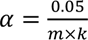.

### Construction of the interactive query system

The system was implemented with the open-source Dash framework, which incorporates the Plotly library to provide dynamic, web-based visualization and user interaction (https://dash.plotly.com/layout).

## Compliance and ethics

### Ethics approval and consent to participate

The Medical Ethics Committee of West China Hospital of Sichuan University approved this study. This study was conducted following the principles of the Helsinki Declaration of 2013.

## Consent for publication

Not applicable.

## Data availability

All data used in this study are provided in the supplementary materials. The all-integrated *H. pylori* genomic resources were submitted to the developed website (https://www.hpgnomad.top/). Reference *H. pylori* strains include *H. cetorium* (https://www.ncbi.nlm.nih.gov/nuccore/CP003479.1/) and *H. pylori* strain 26695 (https://www.ncbi.nlm.nih.gov/nuccore/NC_000915.1?report=genbank). The public data can be downloaded from databases: NCBI (https://www.ncbi.nlm.nih.gov/nuccore/?term=h.pylori), EnteroBase (https://enterobase.warwick.ac.uk/species/helicobacter/search_strains), Datadryad (https://datadryad.org/stash) and PubMLST (https://pubmlst.org/organisms/helicobacter-pylori/).

## Authors’ contributions

G.L.H., M.G.W., R.K.T., H.H.D., B.W.Y., H.L., and L.T.L. conceptualized and supervised the project. L.T.L., B.W.L, J.Z., Y.H.L, H.C., L.M.Z, X.Q.T, Y.T.J., Z.H.Z, Y.Z.W., and Y.G.H conducted data analysis. F.X.B., H.J.Y., C.L., B.W.Y., and H.L. drafted the manuscript, and all the primary authors reviewed, edited, and approved the manuscript.

## Extended Data Figs

**Extended Data Fig. 1.**
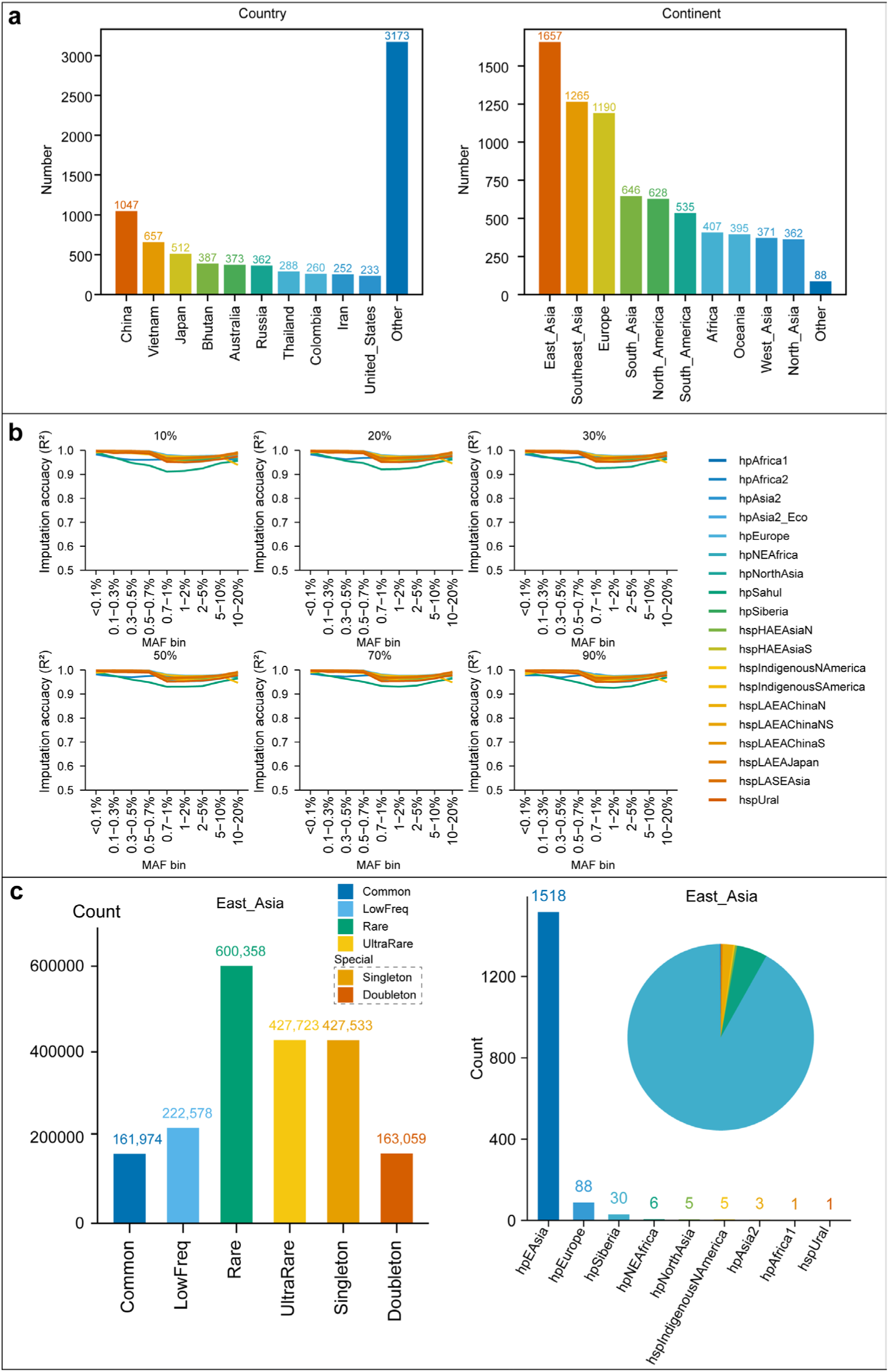
More information of sample distribution in the HPgnomAD database, variant discovery, and reference-panel performance evaluation. **a**, Bar chart of the number of strains collected per country and continent. **b,** Imputation accuracy across coverages, MAF bins, and populations. The x-axis denotes MAF bins, and the y-axis indicates empirical dosage R^2^. **c,** Left: bar chart of variant counts; right: combined bar and pie charts showing population distribution in East Asia.

**Extended Data Fig. 2.**
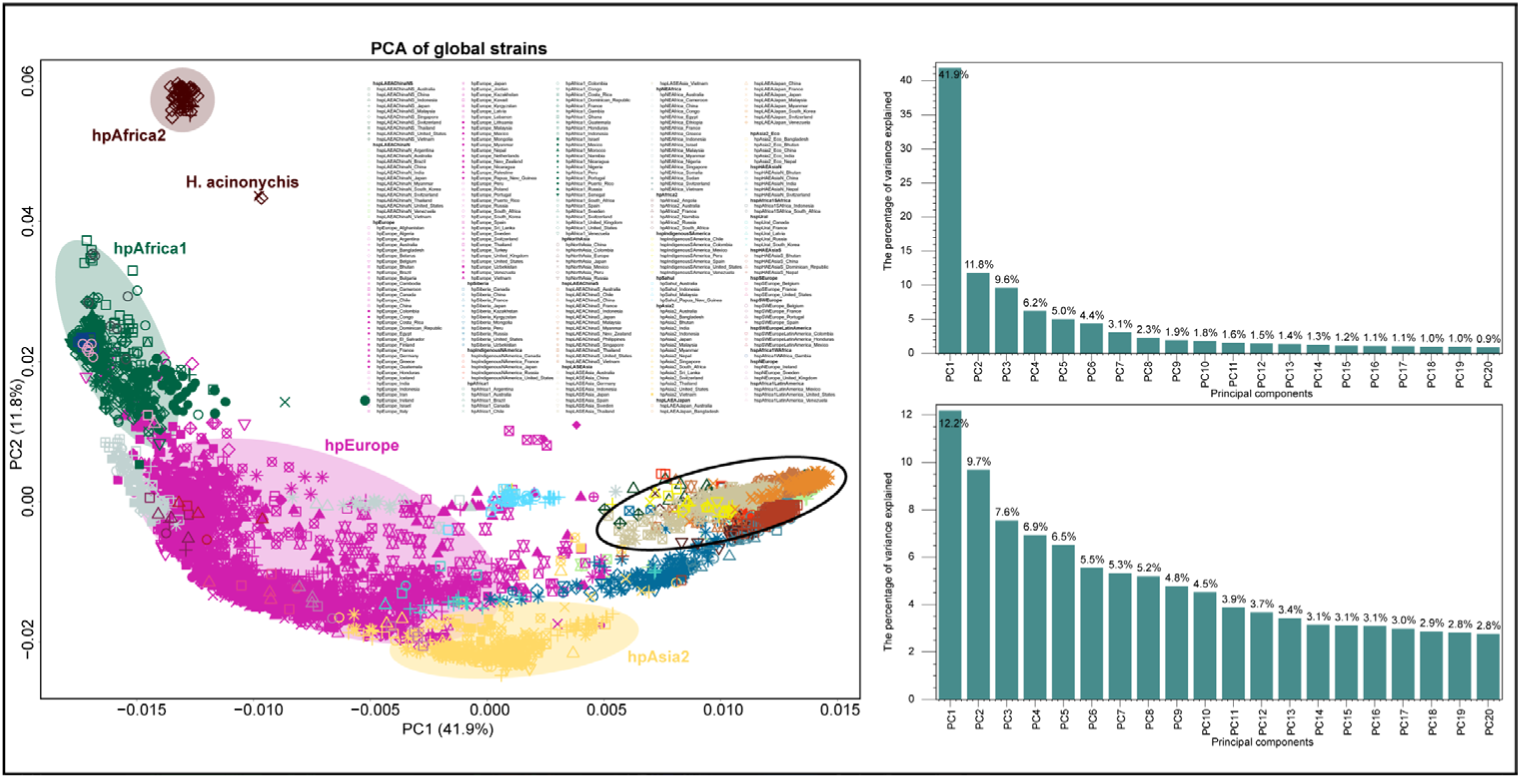
PCA analysis. Left, PCA of global strains, identical to Fig. 2a but colored by genetic background. Upper right, variance explained by the global PCA. Lower right, variance explained by the hpEAsia PCA.

**Extended Data Fig. 3.**
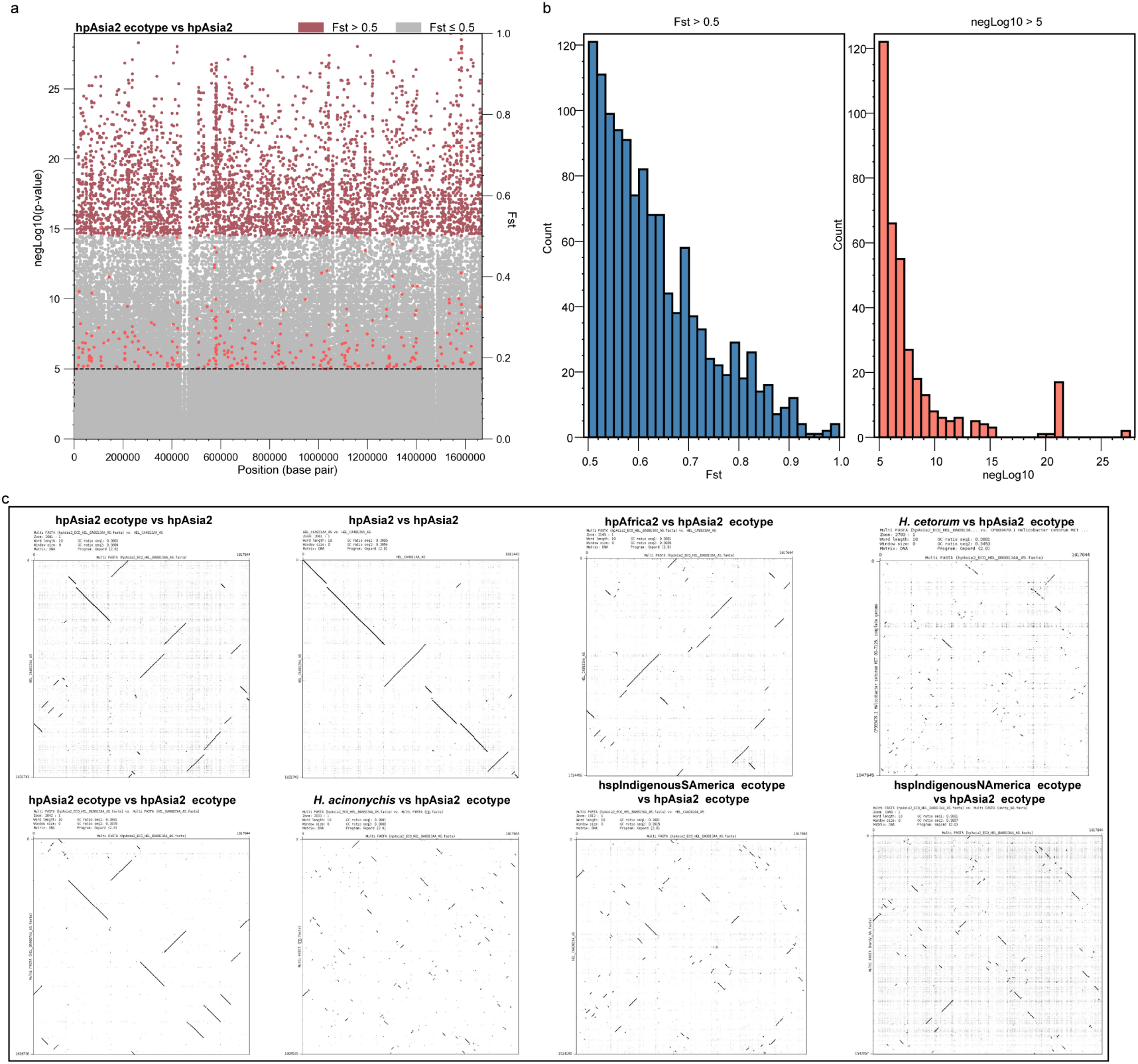
Genomic features of the hpAsia2 ecotype. **a**, Manhattan plot from a GWAS comparing the hpAsia2 ecotype with hpAsia2 strains. Each point represents a SNP; colors denote *F*_ST_ bins (gray and red). The horizontal line marks the genome-wide significance threshold (−log_10_*p* = 5), calculated with a Bayesian Wald test followed by Bonferroni correction (345,807 SNPs tested). **b,** Counts of differentiated loci. Bar charts show the number of sites with *F*_ST_ > 0.5 (left) and with GWAS significance (−log_10_*p* > 5; right). In total, 1,226 high *F*_ST_ loci and 359 GWAS-significant loci were detected, of which 36 satisfy both criteria. **c,** Whole-genome dot-plot comparisons within and between hpAsia2 ecotype genomes (genome pairs indicated above each panel). Continuous diagonals denote collinear genomes; breaks or multiple short diagonals reveal chromosomal rearrangements or differences in gene content, whereas long uninterrupted diagonals indicate highly similar genomes.

**Extended Data Fig. 4.**
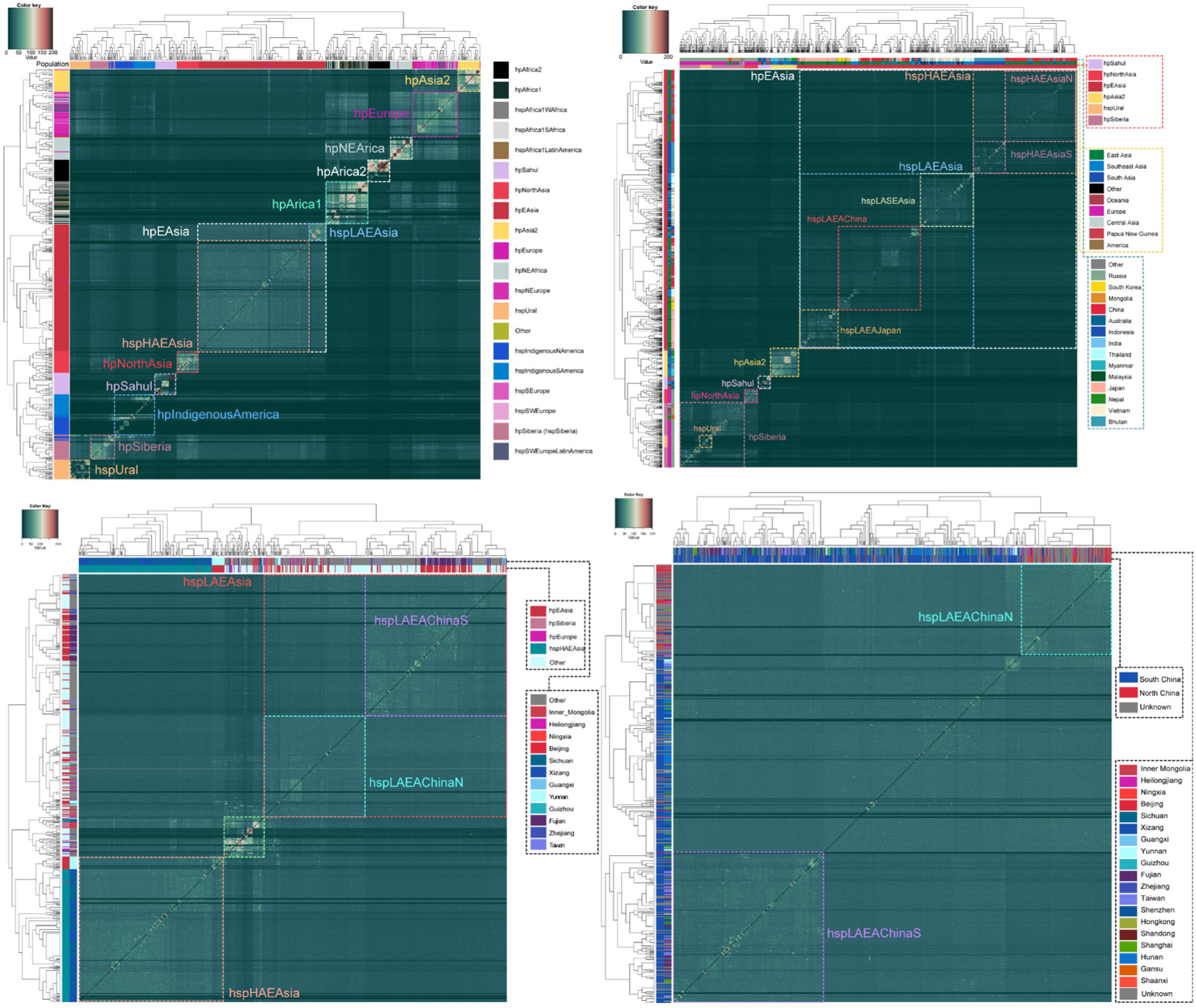
FineSTRUCTURE analysis of strains from global and hpEAsia perspectives. Strain labels defined in previous studies and geographic information are overlaid on the left dendrogram and at the top of the heatmap. Strains that have been redefined or newly defined in this study are highlighted with square markers and bold font. FineSTRUCTURE employs an in-silico chromosome painting algorithm that models each strain as a mosaic of its nearest neighbors, selected from among the other strains in the dataset. Each row in the heatmap represents the coancestry vector for a strain, reflecting the count of DNA segments contributed by each of the other strains. High coancestry between strains indicates that they share numerous genomic segments and likely originate from a common gene pool. Moreover, FineSTRUCTURE-based clustering is more sensitive to recent gene flow compared to traditional clustering methods based on genetic distances.

**Extended Data Fig. 5.**
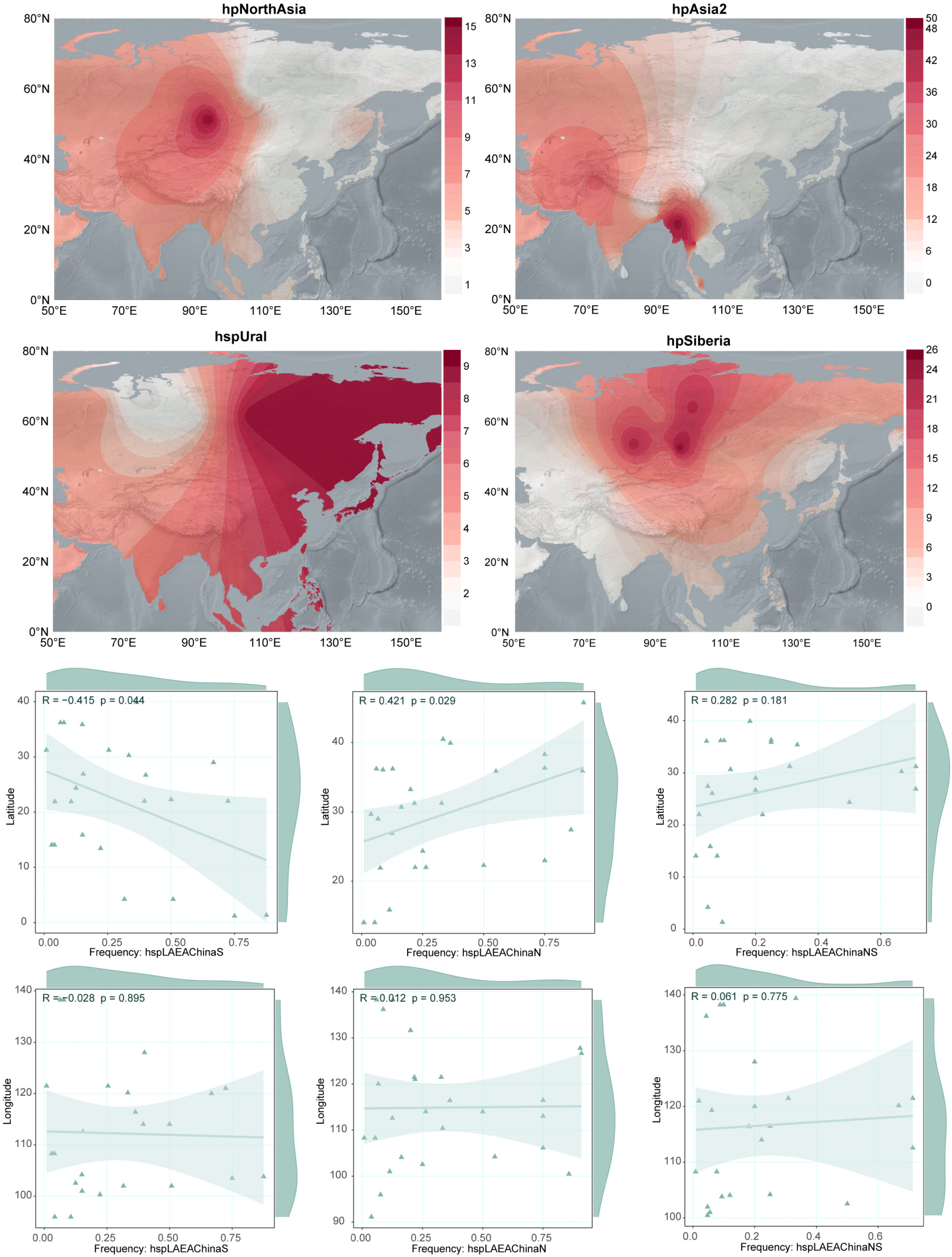
Geographic distribution of H. pylori strains across Asia and correlation analysis of hspLAEAChina. **a**, Spatial distribution patterns of different *H. pylori* strains in Asia are illustrated. Contour maps represent interpolated frequency distributions generated using the Kriging algorithm. **b,** Pearson correlation analysis between latitude, longitude, and frequency of hspLAEAChina strains.

**Extended Data Fig. 6.**
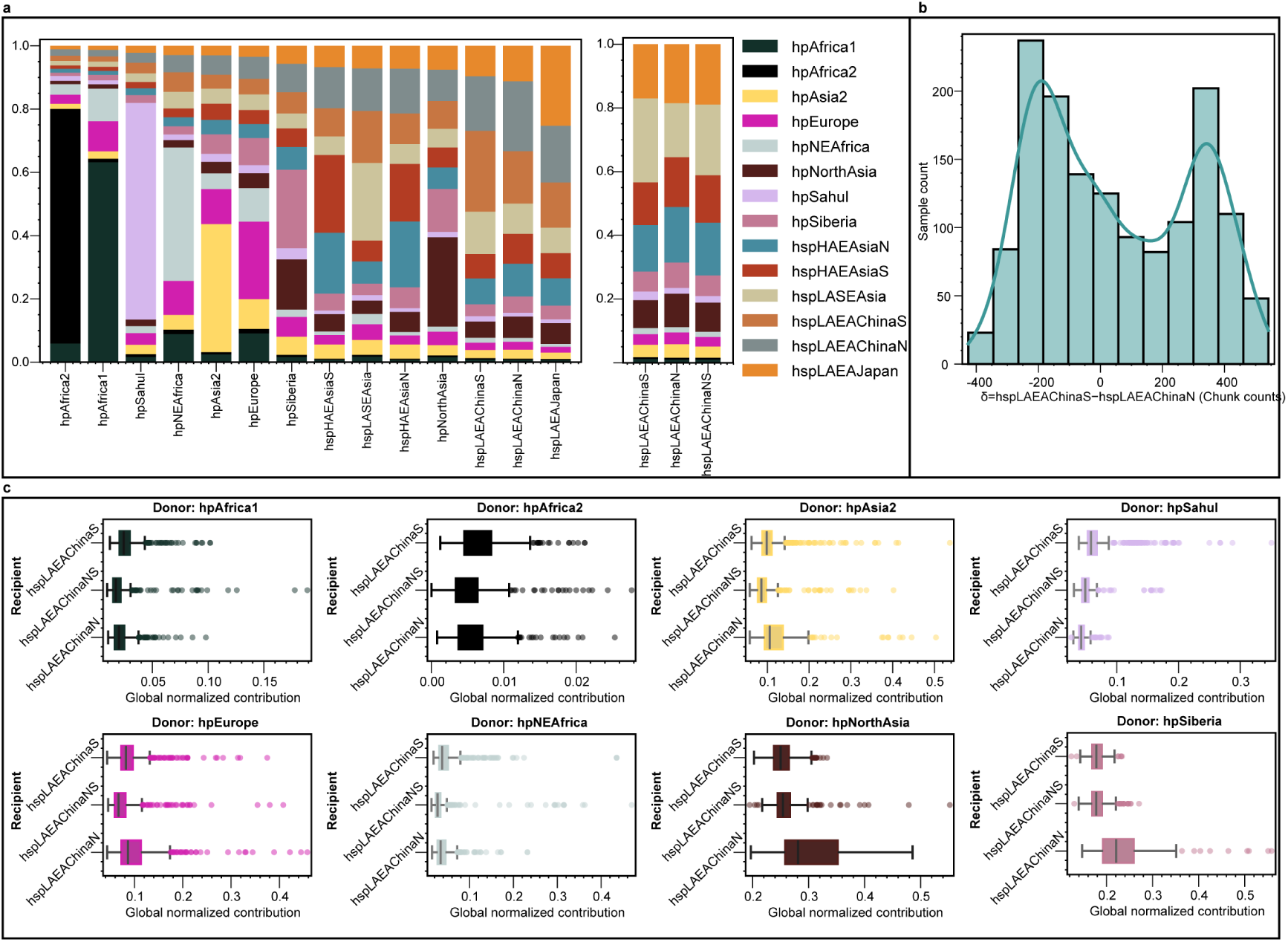
Average ancestry profiles of *H. pylori* in Asia. **a**, ChromoPainter results. The left panel includes the major global strains and all hpEAsia strains as recipients, while the right panel excludes hspLAEAChina from the donor set. **b,** Among the strains assigned to hspLAEAChina, the contributions from hspLAEAChinaS and hspLAEAChinaN differ. **c,** Full version of the box-and-whisker plots illustrating contributions from various donors across the three recipient categories.

**Extended Data Fig. 7.**
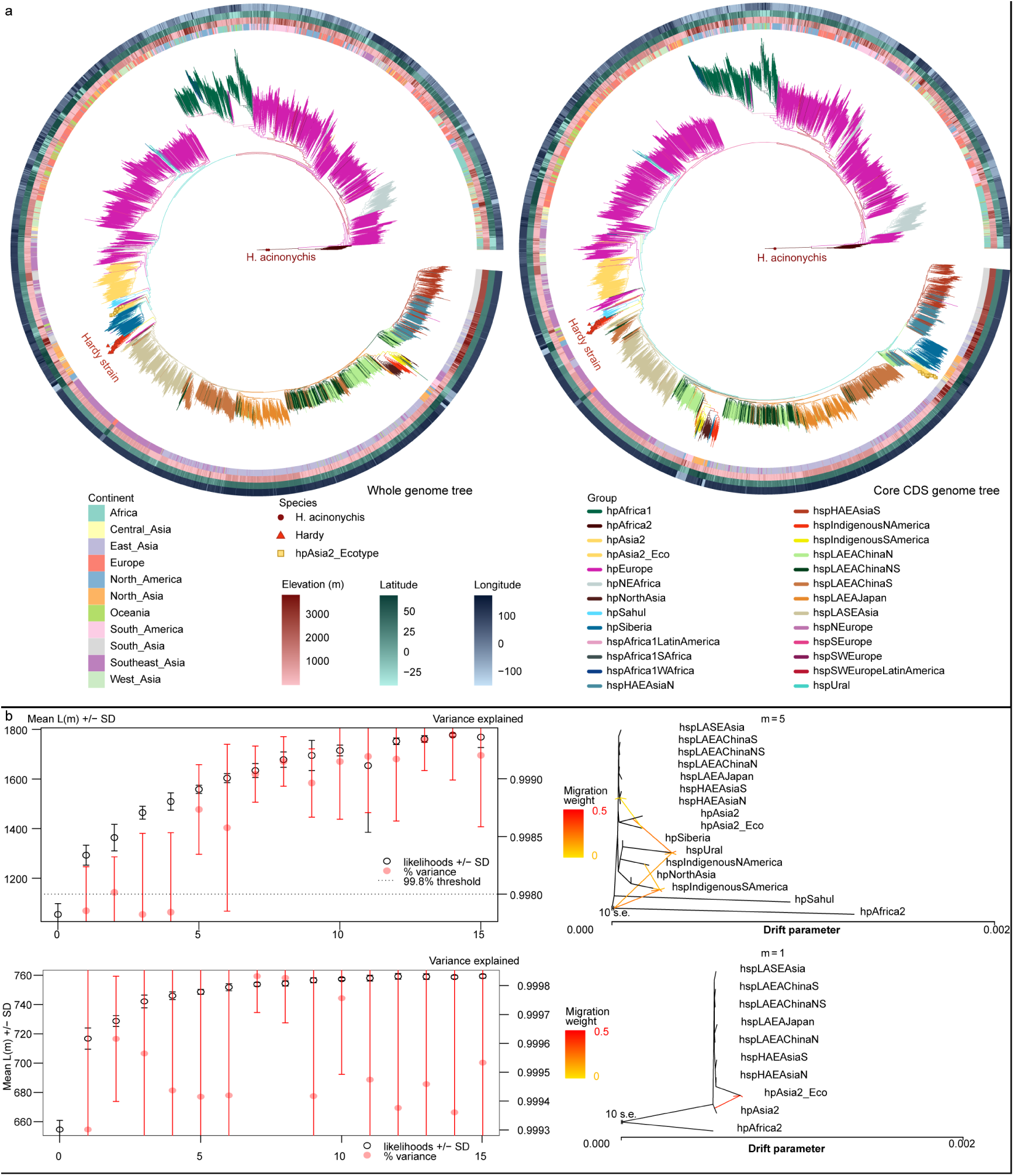
Phylogenetic trees and TreeMix result. **a**, Maximum-likelihood phylogenetic tree based on whole-genome sequences (left) and based on core CDS genome sequences (right). Branches are colored according to population. H. acinonychis serves as the outgroup, indicated by a red square at the tip of its branch. Hardy strains are marked by red triangle dots at the branch tips, and hpAsia2 ecotype are marked by yellow square at the tip of its branch. Concentric circles surrounding each tree represent, from innermost to outermost, the sampling location’s classification by continent, elevation, latitude, and longitude. **b,** Treemix graphs for different populations. The top panel includes strains common across Asia; the bottom panel includes only strains common across East Asia. The optimal number of migration edges was chosen at the inflection point of maximal change in explained variance. The hpAfrica2 population was set as the outgroup. Arrows denote gene flow between branches, with arrow color indicating the migration-edge weight.

**Extended Data Fig. 8.**
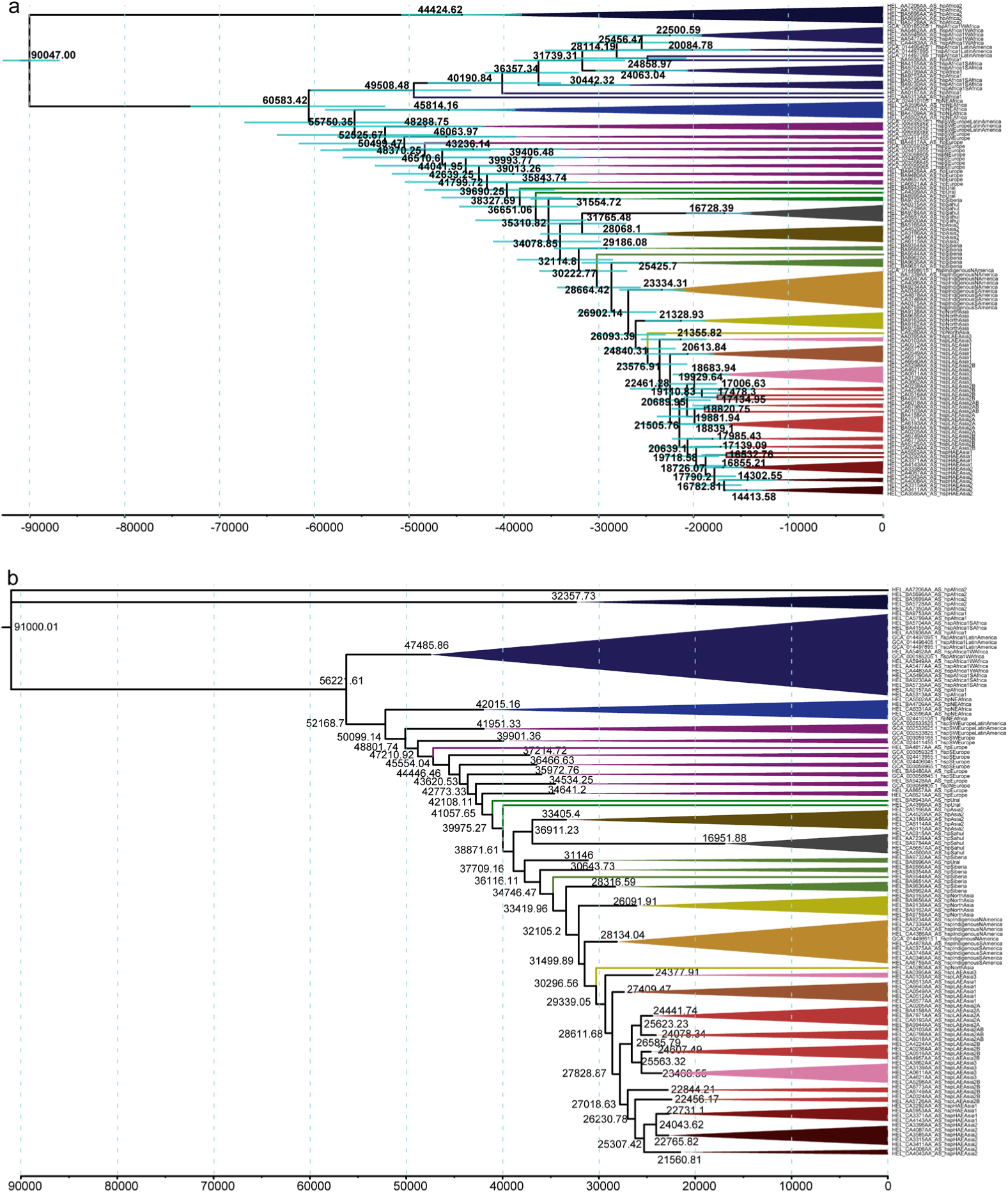
Time scaled phylogeny. **a**, Phylogenetic tree inferred using BEAST, presented here as the complete version of Fig. 4c. **b,** Phylogenetic tree inferred using PAML.

**Extended Data Fig. 9.**
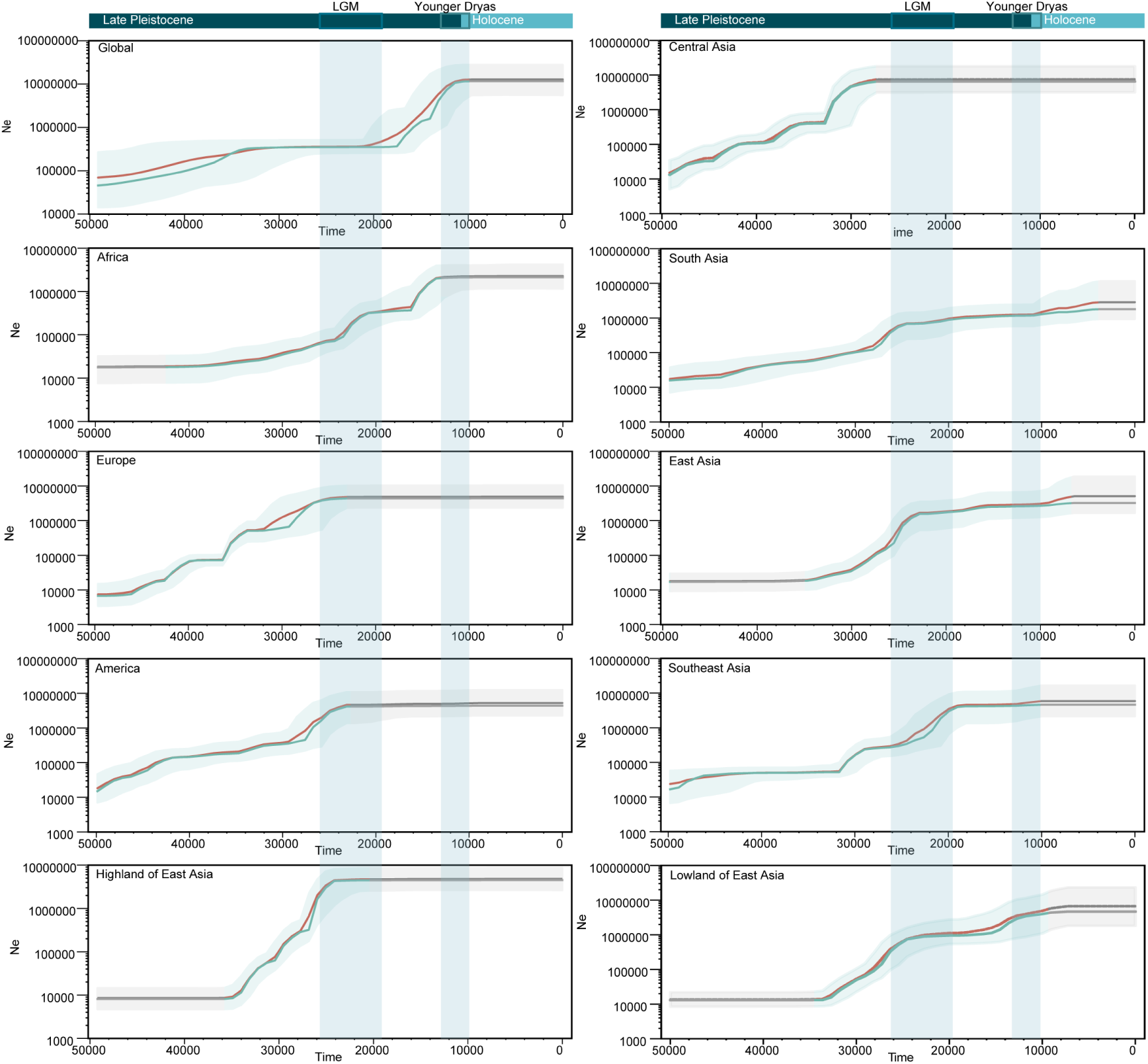
Bayesian skyline plots (BSPs) for each region. The x-axis represents time (in years before the present), and the y-axis denotes effective population size. The shaded area of each BSP curve indicates the minimum-to-maximum range, the green solid line represents the mean value, and the orange dashed line represents the median value. Major geological periods are marked at the top. Grey shaded intervals correspond to periods when the BSP curves plateau, suggesting that genetic data cannot reliably reconstruct population dynamics during those intervals.

**Extended Data Fig. 10.**
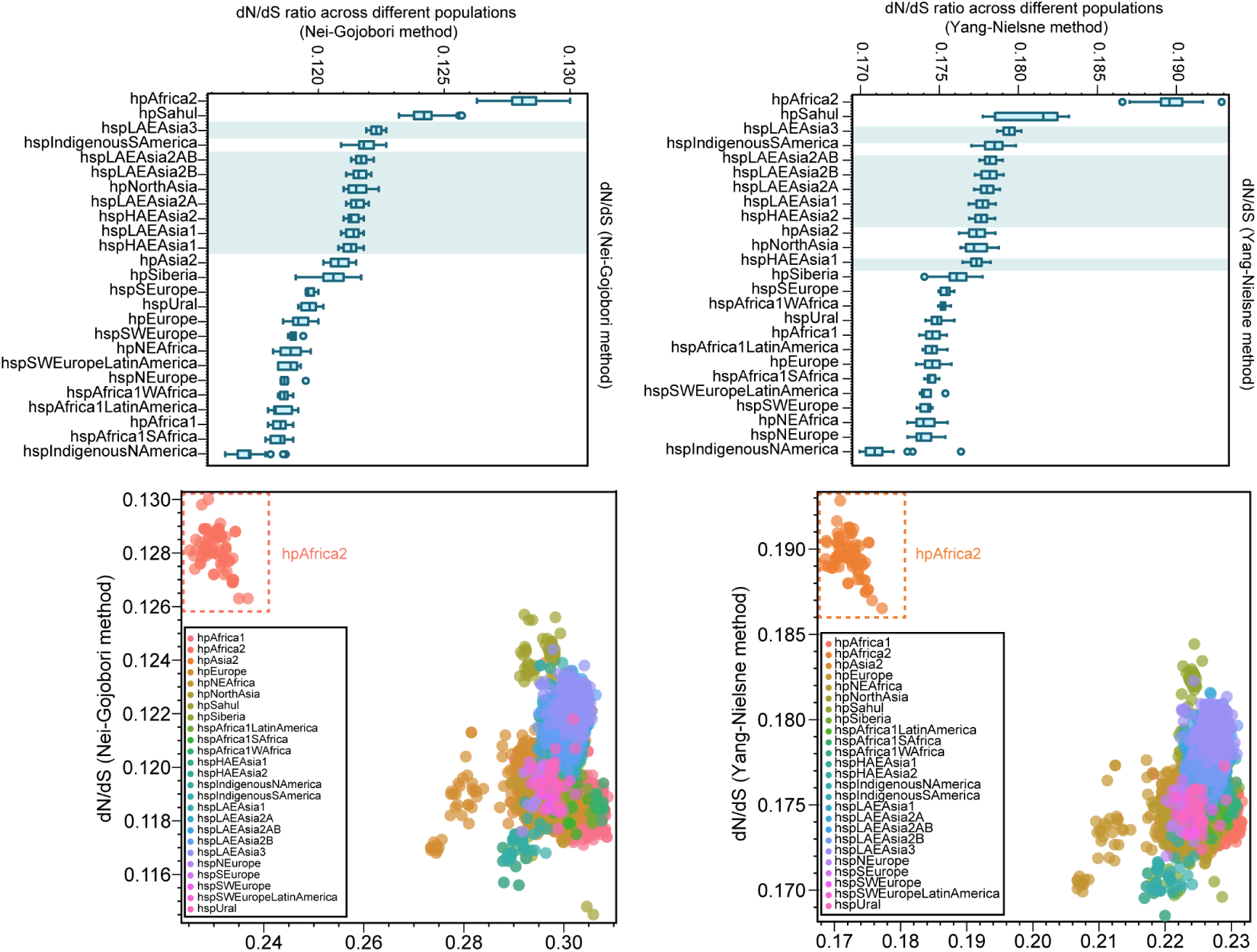
dN/dS estimates for all strains. dN/dS ratios were calculated using the Nei– Gojobori method (left panels) and the Yang–Nielsen method (right panels). The upper panels show box-and-whisker plots summarizing the distribution of *dN/dS* values for each population, whereas the lower panels depict scatter plots of individual gene estimates, with synonymous substitution rates (*dS*) on the x axis and corresponding *dN/dS* ratios on the y axis.

**Extended Data Fig. 11.**
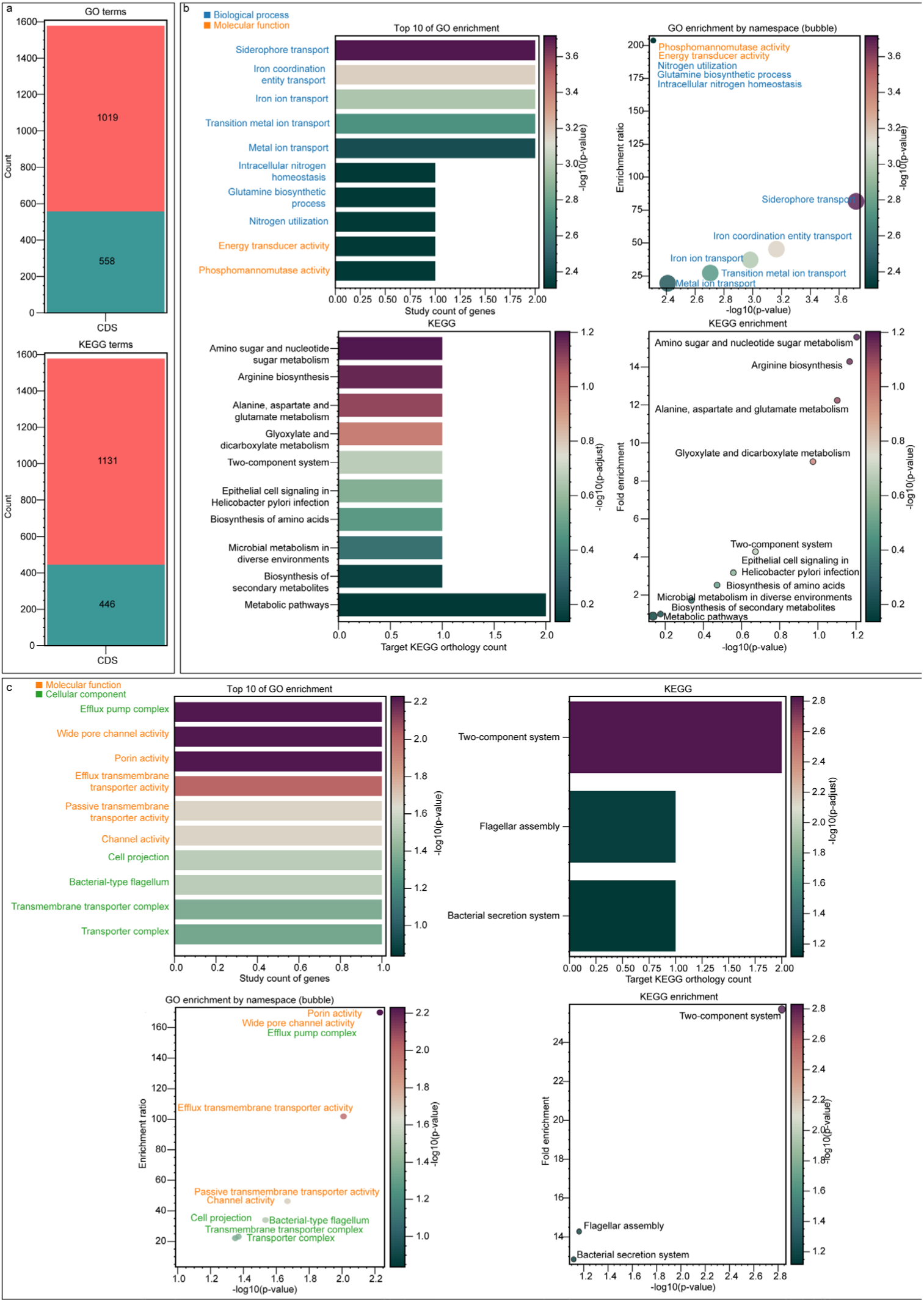
GO and KEGG enrichment analyses. **a**, Bar plot showing, for the 1,577 CDSs, the number of CDSs successfully annotated (red) versus not annotated (green) against GO and KEGG databases. **b,** GO-term and KEGG-pathway enrichment for genes differentiating high-altitude versus low-altitude strains. The target set comprises genes harboring SNPs that satisfy GWAS (−log_10_*p* > 5) or *F*_ST_ > 0.6 thresholds, and exhibit |Cliff’s Δ| ≥ 0.35 in dN/dS comparisons; all other annotated genes serve as the background. Only the top 10 enriched terms are shown. **c,** GO-term and KEGG-pathway enrichment for genes differentiating northern versus southern East Asian strains, using the same target and background sets as in b. Only the top 10 enriched terms are shown.

**Extended Data Fig. 12.**
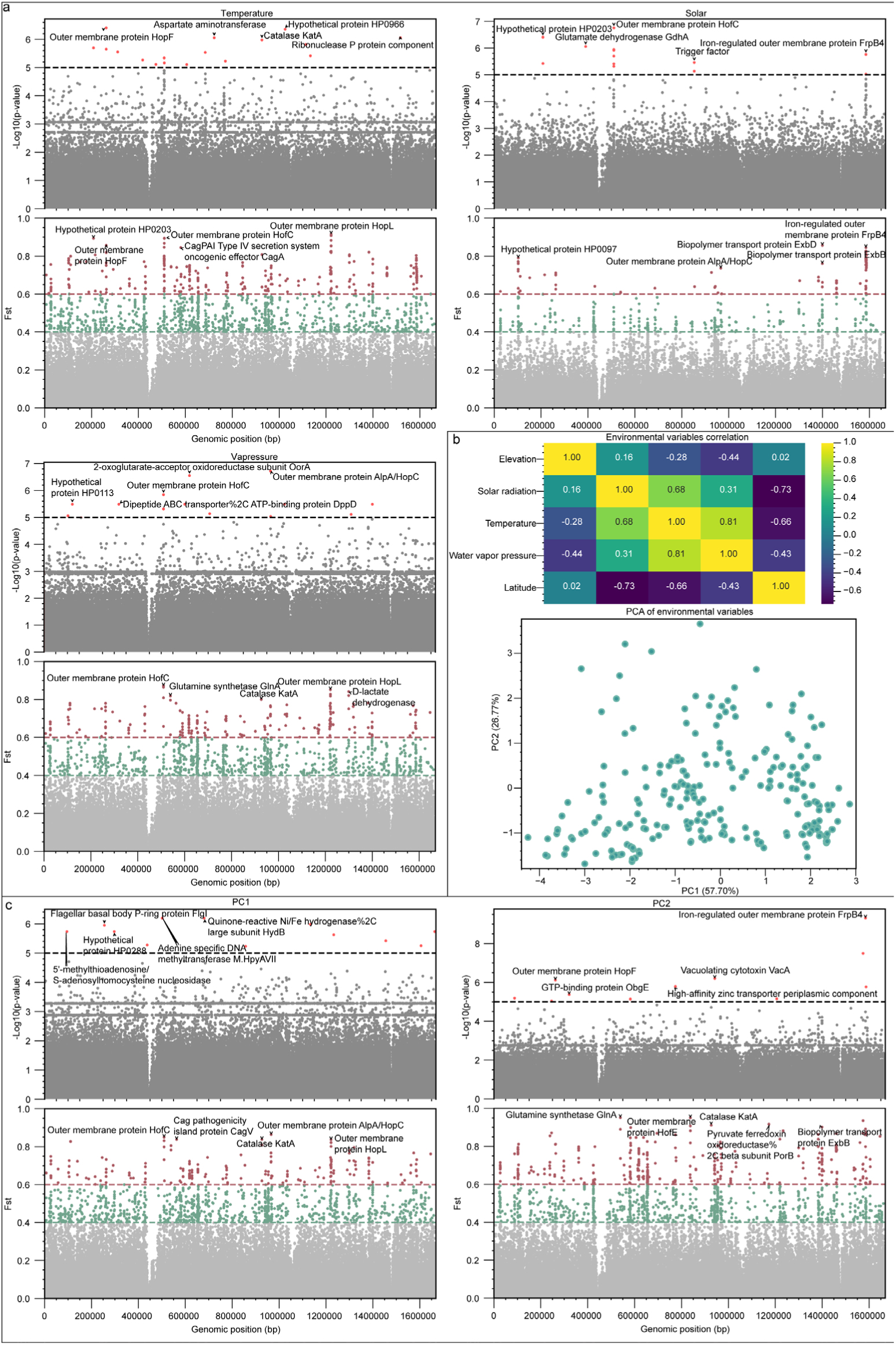
GWAS and F_ST_ analyses for environmental variables. **a**, GWAS and F_ST_ analyses for temperature, solar radiation and vapor pressure. GWAS significance is indicated by a horizontal threshold at −log_10_*p* = 5. SNPs exceeding this threshold are colored red, those below grey. F_ST_ thresholds of 0.4 and 0.6 are marked; SNPs are colored grey (*F*_ST_ < 0.4), green (0.4 ≤ *F*_ST_ < 0.6) or red (Fst ≥ 0.6). **b,** Correlation matrix and PCA of environmental variables. **c,** GWAS and *F*_ST_ analyses for PC1 and PC2 of environmental variables, applying the same significance and *F*_ST_ thresholds as above.

**Extended Data Fig. 13.**
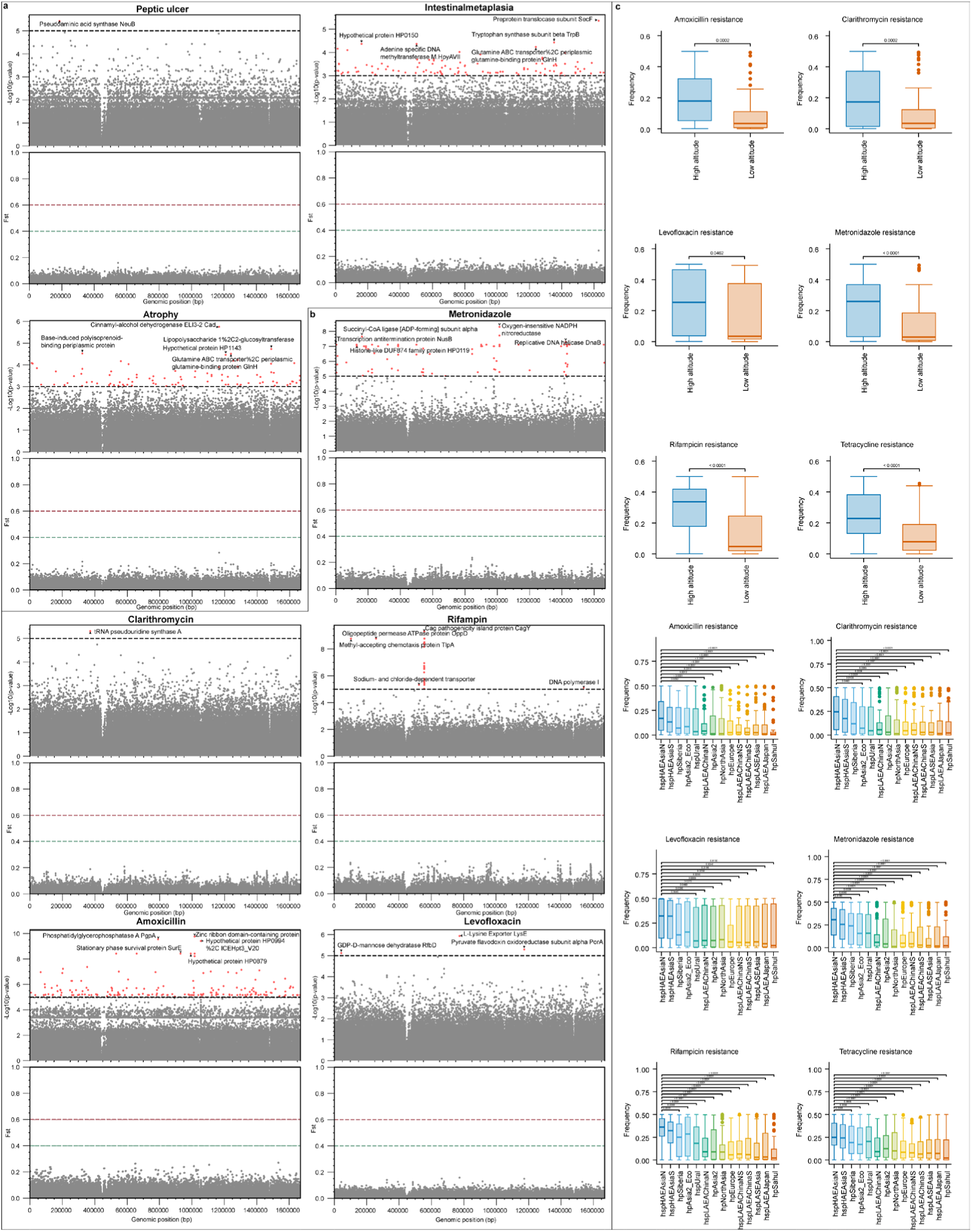
The GWAS and F_ST_ analyses of clinical phenotypes and frequency distribution of candidate loci. **a**, GWAS and F_ST_ analyses of host clinical traits. Panels show GWAS and F_ST_ results for peptic ulcer, intestinal metaplasia and gastric atrophy. Because the sample sizes for intestinal metaplasia and atrophy were limited (n = 133), a relaxed significance threshold of –log_10_ (*p*-value) = 3 was applied. **b,** GWAS and *F*_ST_ results for bacterial antibiotic-resistance phenotypes. Analyses are presented for metronidazole, clarithromycin, rifampicin, amoxicillin and levofloxacin. **c,** Boxplots showing the allele frequency distribution of candidate loci across populations. Candidate loci were those in the top 5% of each analysis based on empirical thresholds, intersected with the top 5% of loci from the high-versus-low altitude differentiation scan. Allele frequencies were computed for each locus in each population.

**Extended Data Fig. 14.**
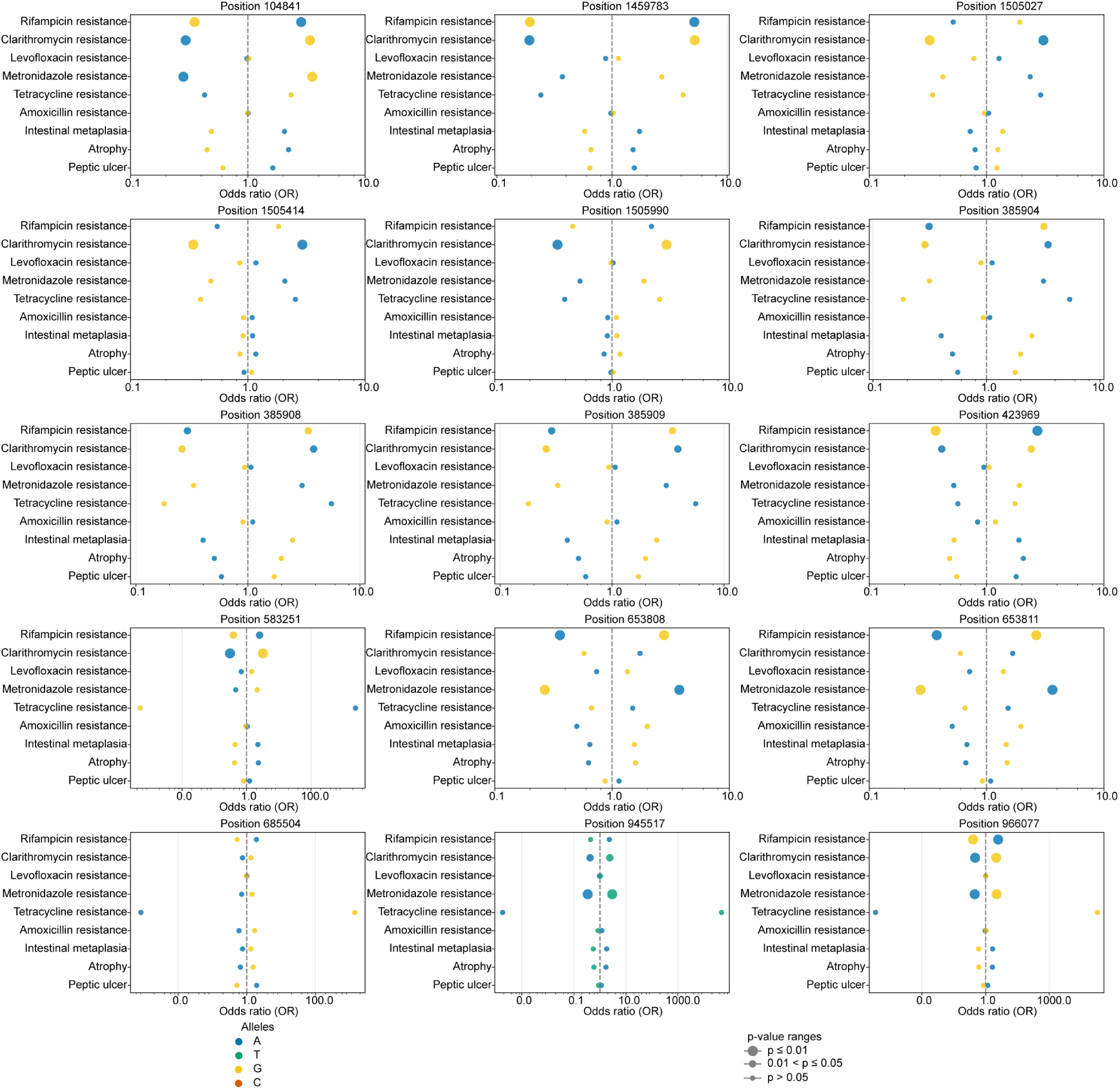
Bubble plot of odds ratios for pleiotropic SNPs. The x-axis shows ORs on a log scale; the y-axis lists phenotypes. Bubble color indicates allele variant, and bubble size reflects Bonferroni-adjusted *p*-value categories (*p* ≤ 0.01; 0.01 < *p* ≤ 0.05; *p* > 0.05). The dashed vertical line marks OR = 1. ORs and *p*-values were derived from logistic regression with Wald tests and Bonferroni correction. This panel presents the full loci including Fig. 6.

